# The plant nuclear lamina disassembles to regulate genome folding in stress conditions

**DOI:** 10.1101/2022.12.01.517823

**Authors:** Nan Wang, Zhidan Wang, Sofia Tzourtzou, Xu Wang, Xiuli Bi, Julia Leimeister, Linhao Xu, Takuya Sakamoto, Sachihiro Matsunaga, Andreas Schaller, Hua Jiang, Chang Liu

## Abstract

The nuclear lamina (NL) is a complex network of nuclear lamins and lamin-associated nuclear membrane proteins, which scaffold the nucleus to maintain structural integrity. In *Arabidopsis* thaliana, Nuclear Matrix Constituent Proteins (NMCPs) are essential components of the NL and are required to maintain the structural integrity of the nucleus and specific perinuclear chromatin anchoring. At the nuclear periphery, suppressed chromatin overlapping with repetitive sequences and inactive protein coding genes are enriched. At a chromosomal level, plant chromatin organization in interphase nuclei displays flexibilities in response to various developmental cues and environmental stimuli. Based on these observations in *Arabidopsis*, and given the role of *AtNMCP* genes (*CRWN1* and *CRWN4*) in organizing chromatin positioning at the nuclear periphery, one can expect considerable changes in chromatin-NL interactions when the global chromatin organization patterns are being altered in plants. Here, we report the highly flexible nature of plant nuclear lamina, which disassembles substantially under various stress conditions. Particularly, under heat stress, we reveal that chromatin domains, initially tethered to the nuclear envelope, remain largely associated with CRWN1 and become scattered in the inner nuclear space. Via investigating the three-dimensional chromatin contact network, we further reveal that CRWN1 proteins play a structural role in shaping the changes in genome folding under heat stress. Also, CRWN1 acts as a negative transcriptional co-regulator to modulate the shift of the plant transcriptome profile in response to heat stress.

## Introduction

The nuclear lamina is a layer of protein meshwork underneath the nuclear envelope, in which nuclear lamin proteins are the main constituents. Besides providing mechanical support to the nucleus, the nuclear lamina also participates in regulating many events in the nucleus, such as gene expression, chromatin organization, DNA repair, and DNA replication (Gruenbaum et al. 2005). The plant nuclear lamina is composed of a group of plant-specific coiled-coil domain-containing proteins and lamin-binding membrane proteins, among which those belonging to the Nuclear Matrix Constituent Proteins (NMCPs) family were first discovered and have been investigated most extensively (Ciska et al. 2019; Groves et al. 2020). NMCPs are highly conserved in the plant kingdom, but they do not share any sequence homology with animal nuclear lamins. Amongst the NMCP(s) in each plant species that have been studied so far, there is at least one showing preferential localization at the nuclear periphery (Dittmer et al. 2007; Kimura et al. 2010; Ciska et al. 2013; Sakamoto and Takagi 2013; Ciska et al. 2018; Ciska et al. 2019; Yang et al. 2020; McKenna et al. 2021; Wang et al. 2021). In the basal land plant *Marchantia*, its sole NMCP homolog appears dispensable for vegetative growth (Wang et al. 2021); however, in the higher plant *Arabidopsis*, its NMCP genes are essential to plant viability (Wang et al. 2013). Genetic cross of mutants of different *Arabidopsis NMCP* genes, which are also named *CRWNs (CROWDED NUCLEI*), has revealed functional redundancy of *CRWN* genes. In an earlier study by Wang and colleagues, it was found that quadruple *crwn* mutants, as well as some triple *crwn* mutants, could not be recovered from a segregating population, indicating the requirement of a minimum level of CRWN activities to complete plant life cycle (Wang et al. 2013).

*Arabidopsis* CRWN1 and CRWN4 proteins are located at the nuclear periphery; mutant plants of these *CRWN* genes develop spherical nuclei, whose sizes are also noticeably smaller than those in wild-type plants (Dittmer et al. 2007; Sakamoto and Takagi 2013). On the contrary, overexpression of *CRWN1* promotes nuclear deformation, leading to the formation of ring-like and bleb-like structures on the nuclear envelope (Goto et al. 2014). Transcriptomic analyses of their single and double mutants revealed that the gene expression profile of these mutants was featured with ectopic activation of pathways related to stress responses (Choi et al. 2019). Several transcription factors and co-regulator were identified as interacting partners of CRWN1, suggesting the involvement of plant nuclear lamina in regulating gene expression (Pawar et al. 2016; Guo et al. 2017). Reminiscent of the *crwn* mutant transcriptome, mutants of lamin-interacting membrane proteins in the PNET2 family also show activated stress pathways (Tang et al. 2022).

The *Arabidopsis* nuclear lamina selectively interacts with chromatin regions, creating a non-random chromatin distribution pattern at the nuclear periphery where repressed chromatin is enriched (Bi et al. 2017; Hu et al. 2019). Recent work showed that CRWN1 and CRWN4 help in establishing a scattered distribution of centromeres in the nucleus (Sakamoto et al. 2022). At a chromosomal level, plant chromatin organization in interphase nuclei displays flexibilities. Many developmental cues and environmental factors, such as dedifferentiation (Tessadori et al. 2007a), leaf development (Bourbousse et al. 2015), seedling growth (Mathieu et al. 2003), floral transition (Tessadori et al. 2007b), seed development (van Zanten et al. 2011), light intensity (Barneche et al. 2014; Bourbousse et al. 2015), microbe infection (Pavet et al. 2006), and temperature stress (Pecinka et al. 2010), can trigger global rearrangement of chromatin, demonstrating a tight connection between the structural arrangement of chromatin and its activities. Besides, scattered evidence implies that the plant nuclear lamina might undergo active turnover (Guo et al. 2017; Huang et al. 2020). Thus, given the role of *NMCP* genes (i.e., *CRWN1* and *CRWN4*) in organizing chromatin at the nuclear periphery (Hu et al. 2019), it remains unclear whether the chromatin-NP interactions are dynamic when the global chromatin organization patterns are being altered in plants.

In this study, we show that plant nuclear lamina disassembles substantially following several types of abiotic stress treatment including heat stress. During heat stress response, chromatin domains, initially tethered to the nuclear envelope, remained largely occupied by CRWN1 and become scattered in the inner nuclear space. We further show that during heat stress, CRWN1 proteins function as structural factors to guide genome to adapt an alternative folding conformation. Meanwhile, CRWN1 also acts as a negative transcriptional co-regulator to modulate the shift of the plant’s transcriptome profile in response to heat stress.

## Results

### Changes in chromatin positioning under heat stress

Our recent work on CRWN1 and CRWN4 indicated that they were required in establishing specific perinuclear chromatin localization patterns (Hu et al. 2019). At the nuclear periphery, CRWN1 proteins preferentially interact with chromatin regions having low accessibility, which overlap with inactive protein-coding genes and transposons (Hu et al. 2019). Given the fact that plants have extensive chromatin-based transcriptional reprogramming during stress responses (Bhadouriya et al. 2020), we asked whether stress could trigger changes in the chromatin-nuclear lamina interaction patterns. Of many widely studied abiotic stress conditions, we chose heat stress in our initial experiment because of the availability of published data documenting changes in transcriptome, chromatin structure, and genome organization during heat stress adaptation (Pecinka et al. 2010; Sun et al. 2020), which together implied possible dynamic chromatin positioning at the nuclear periphery. To this end, we selected a few genomic loci belonging to nuclear-lamina-associated domains (LADs) and examined their localization in heat-stressed and control plants (Figure 1A). In control plants (i.e., without heat stress treatment), FISH (fluorescent *in situ* hybridization) probes recognizing LAD loci showed preferential localization near the nuclear envelop; on the contrary, such perinuclear positioning was lost in heat-stressed plants (Figure 1B and 1C). Besides, our FISH experiment revealed a higher incidence of chromatin decondensation of the probed genomic regions in stress-treated plants, which was similar to the changes in centromeric repeats triggered by heat (Pecinka et al. 2010)(Figure S1). This observation agrees with a recent study showing global rearrangement of chromatin organization induced by heat (Sun et al. 2020), and further implies that it involves the detachment of LADs from the nuclear envelope. To address whether LADs lost their contacts to the nuclear lamina under heat stress, we performed chromatin immunoprecipitation (ChIP) assays to examine the interactions between LAD loci and CRWN1. The CRWN1 ChIP was performed with a *CRWN1:2HA* tagging line, in which the fusion protein between CRWN1 and a tandem HA (Hemagglutinin) tag could fully rescue *crwn1* loss-of-function phenotypes (Hu et al. 2019). We found enrichment of the selected LAD loci by CRWN1:2HA in both control and heat-stressed plants; to our surprise, heat treatment did not weaken the interactions between CRWN1 and these loci (Figure 1D). Since LAD loci lost their perinuclear localization in heat-stressed plants, we therefore proposed that the CRWN1 protein, and perhaps other components of the nuclear lamina, acquired alternative nuclear localization patterns.

**Figure 1.**
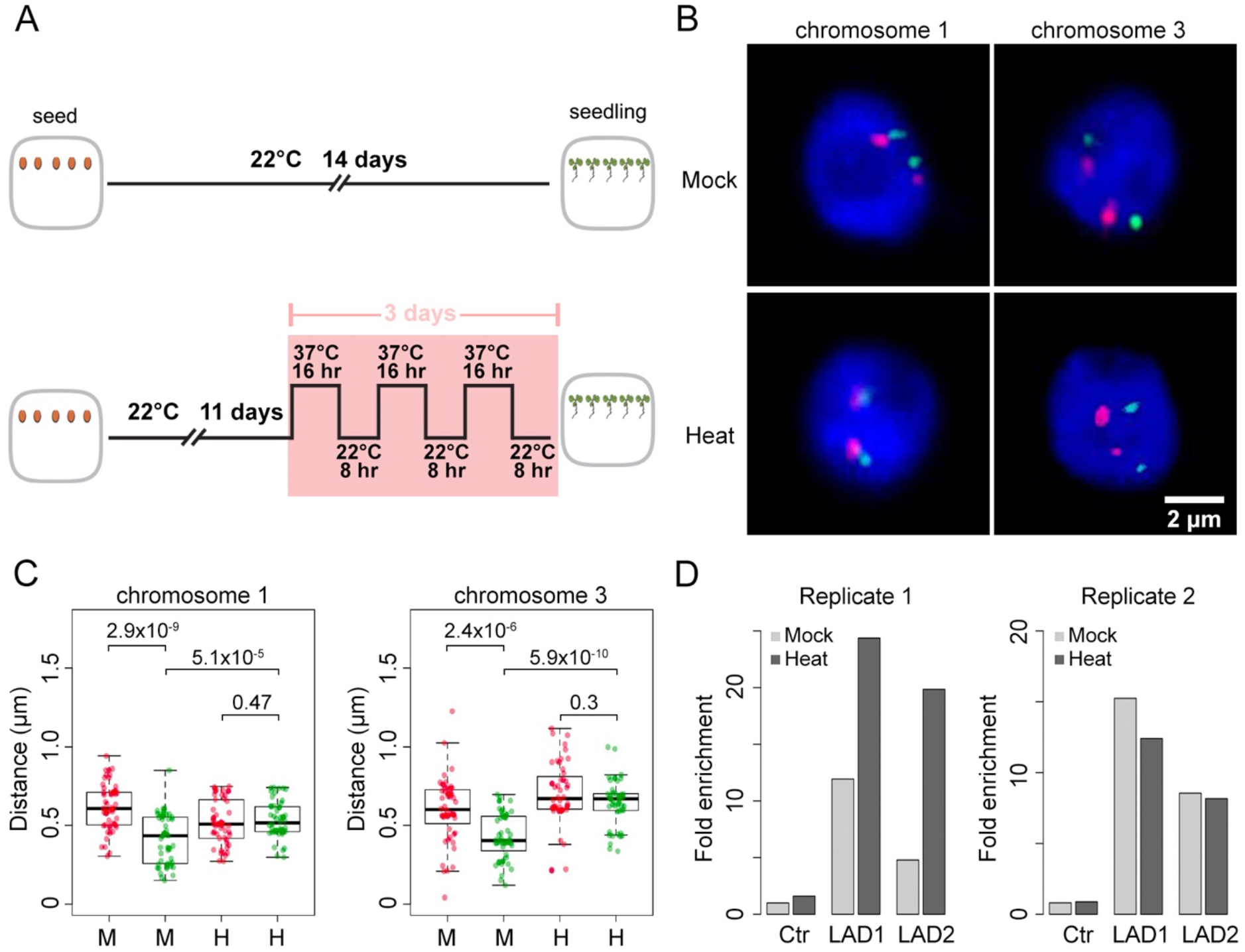
LAD chromatin detaches from the nuclear periphery under heat stress. (A) The experiment setup. (B and C) Representative FISH images and quantification showing the localization of LAD (green) and non-LAD chromatin with respect to the nuclear periphery. The p-values shown above the boxplots were obtained from twosided Mann-Whitney U tests. M and H stand for mock- and heat-stress treatment, respectively. (D) ChIP-qPCR showing the interactions between CRWN1 and two LAD-loci, which overlap with gene loci *AT1G65750* and *AT3G29767*, respectively. “Ctr” refers to a non-LAD locus. The relative fold enrichment was calculated by using the *TUB2* genomic locus as the reference.

### Responses of the plant lamina to stress conditions

The plant nuclear lamina is a filamentous network mainly built up with NMCPs (Masuda et al. 2021). In *Arabidopsis*, two NMCP proteins, CRWN1 and CRWN4, are known to be involved in establishing such a protein meshwork (Sakamoto and Takagi 2013). Another protein named KAKU4 is also a constitutive member of the *Arabidopsis* nuclear lamina (Goto et al. 2014). To address whether the nuclear lamina acquires an alternative organization under heat stress, we examined its members’ protein localization patterns. In total, we analyzed CRWN1:2HA, CRWN4:2HA, and KAKU4:GFP (Goto et al. 2014). We also examined proteins that were located at the nuclear periphery but were not directly involved in assembling the nuclear lamina. The selected proteins were NUP1:GFP, which was a nuclear basket protein belonging to the nuclear pore complex, and SUN1:GFP, which was an integral protein located in the inner nuclear membrane (Ostlund et al. 2009; Tamura et al. 2010; Pawar et al. 2016). We found that nuclear lamina proteins (i.e., CRWN1, CRWN4 and KAKU4) lost their specific perinuclear location during heat stress; whereas SUN1 and NUP1 did not (Figure 2A-C, Figure S2). Besides, we did not find any protein degradation or cleavage associated with these nuclear lamina component proteins (Figure 2D). In fact, CRWN4 appeared with higher abundance after heat treatment (Figure 2D). These findings indicate that during heat stress response, the nuclear lamina disassembles, along which its multiple components acquire localization in the nuclear interior.

**Figure 2.**
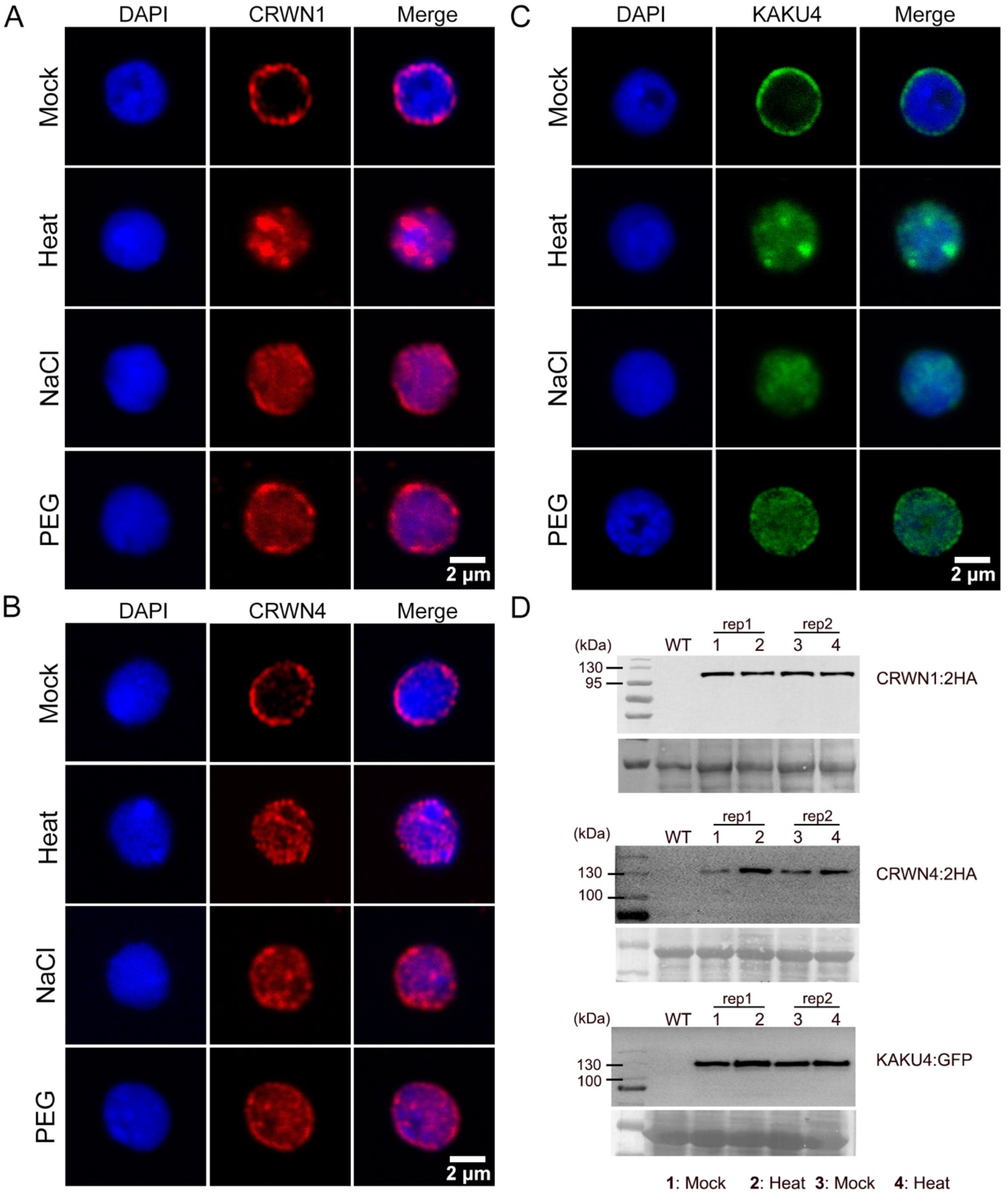
*Arabidopsis* nuclear lamina component proteins lost preferential perinuclear localization under abiotic stresses. (A-C) Each panel shows representative confocal images of the central focal plane of nuclei. The changes in protein localization of CRWN1 (A), CRWN4 (B), and KAKU4 (C) in response to various abiotic stresses was observed in all randomly selected nuclei (n > 15) in two independent batches of experiments. “NaCl” and “PEG” refer to treating plant with 150 mM NaCl and 5% Polyethylene Glycol for two days, respectively. (D) Plant nuclear lamina components are not cleaved or degraded under heat stress. The photos shown blew individual immunoblots depict loading control. WT indicates wild-type protein extract.

Previous functional studies on *Arabidopsis* nuclear lamin component genes indicated that they participated in regulating nuclear size and shape (Dittmer et al. 2007; Sakamoto and Takagi 2013; Goto et al. 2014). We asked if the disassembly of nuclear lamina during heat shock could cause changes in nuclear morphology, making them resemble nuclei in lamin mutant plants. Compared to control plants, heat-stressed plants surprisingly maintained nuclear shape and size (Figure S3). In addition, our statistical analyses on nuclear sizes revealed a clear difference in nuclear morphology between heat-stressed wild-type and *crwn1* plants, indicating that nuclear lamin proteins, albeit having distinct nuclear localization under heat stress, are indispensable to maintain nuclear structure (Figure S3).

We further asked if other stress conditions were able to trigger similar changes to the nuclear lamina. By treating plants with salt or osmotic stresses, we also observed remarkable detachment of CRWN1, CRWN4 and KAKU4 from the nuclear periphery (Figure 2). On the contrary, SUN1 and NUP1 remained their perinuclear localization under these stress conditions (Figure S2). Interestingly, after plants have been exposed to a stress condition for a long time, e.g., seedlings continuously facing 50 mM NaCl stress from starting germination, CRWN1 protein abundance largely dropped and the remaining proteins were distributed throughout the nucleoplasm (Figure S4). Taken together, these results reveal the dynamic nature of the *Arabidopsis* nuclear lamina; upon perceiving different stress conditions, it disassembles and releases its component proteins into the nuclear interior.

### Dynamic lamin-chromatin interactions under heat stress

Following our findings that LAD chromatin remained in direct contact with CRWN1 proteins after the disassembly of nuclear lamina in heat response (Figure 1 and Figure 2), we sought to gain a better view on the global interactions between chromatin and CRWNs in this process. To this end, we performed ChIP-seq experiments by using the *CRWN1:2HA* and a newly created *CRWN4:2HA* tagging lines in their respective mutant genetic backgrounds. The CRWN4:2HA tagging line carried a genomic *CRWN4* sequence fused with tandem HA repeats, which could fully rescue the nuclear morphology phenotypes (smaller and round nuclei) in *crwn4* (Figure S5). Principal component analysis of ChIP-seq sample reads distribution patterns revealed noticeable changes in CRWN1-chromatin interactions before and after heat stress but not in CRWN4:2HA-chromatin interactions (Figure S6). Besides, close examination of ChIP-seq signals across individual chromosomes indicated that among CRWN1 and CRWN4, the former was the overwhelming protein interacting with chromatin (Figure 3A and Figure S7). The majority of CRWN4:2HA-bound chromatin, which was also enriched with CRWN1:2HA, was located in centromeric and pericentromeric regions (i.e., chromocenter) (Figure 3B). We also observed that heat stress influenced CRWN1- and CRWN4-chromatin interactions differentially. Compared to control plants, heat-stressed plants showed stronger CRWN1:2HA-chromatin interactions in chromocenter regions, but the interactions in chromosome arms became weaker (Figure 3C and Figure S7C). Such a shift of CRWN1:2HA-chromatin interaction patterns suggests that decondensed chromocenters, triggered by heat-stress, promotes the formation of new contacts with CRWN1:2HA. Although similar amount of genomic regions were enriched with CRWN1 before and after heat stress (23.26 Mb in control vs. 21.72 Mb under heat), they only shared 15.42 Mb (Figure 3C). On the contrary, heat did not result in a shift of CRWN4:2HA-chromatin interaction patterns but in general it made them weaker. Of the 8.6 Mb CRWN4:2HA-enriched regions in control plants, 4.27 Mb (50%) were lost in heat-stressed plants; while those regions that gained enrichment under heat stress were only 0.88 Mb (Figure 3C). Interestingly, CRWN4:2HA-chromatin interactions were stronger at the boundary of CRWN1:2HA-enriched regions, implying that CRWN1 and CRWN4 may occupy different space at the interface of the plant lamina and chromatin (Figure 3D). In support to this notion, a recent study employing high-resolution microscopy revealed that NMCP1 and NMCP2 proteins in *Apium graveolens* form distinct networks at the nuclear periphery (Masuda et al. 2021). Altogether, we conclude that *Arabidopsis* CRWN1 is the main NMCP protein involved in forming nuclear lamina-chromatin contacts at the nuclear periphery, and these contacts largely remain under heat stressed condition along with the disassembly of the nuclear lamina.

**Figure 3.**
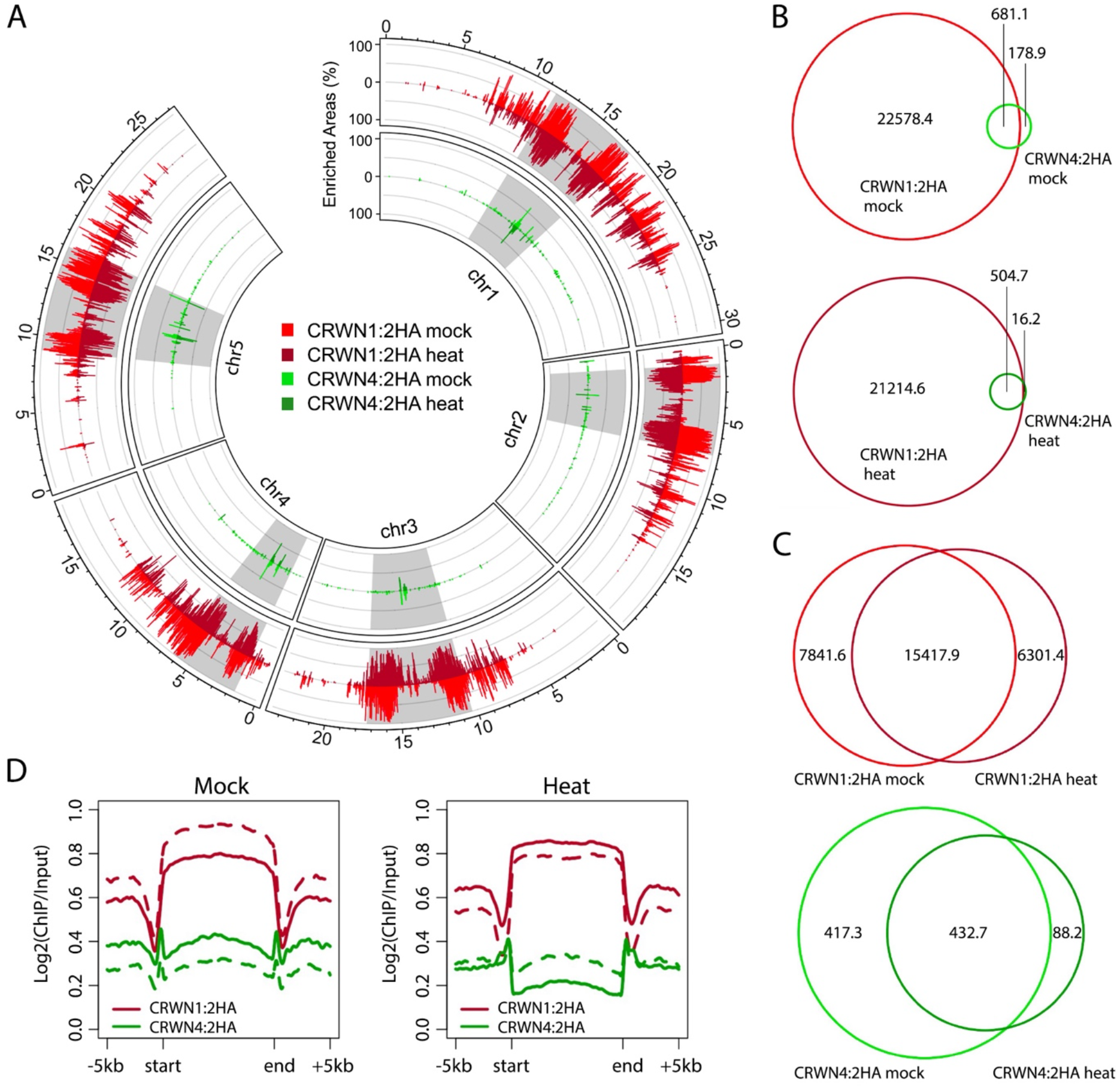
A global view of CRWN-chromatin association under heat stress. (A) A genomic overview of CRWN1- and CRWN4-enriched chromatin. The gray blocks depict pericentromeric regions. (B,C) Venn Diagram of genomic regions (unit: kb) enriched in individual samples. (D) ChIP-seq signals of CRWN1 and CRWN4 proteins across CRWN1-enriched genomic regions. The solid and dashed lines depict two biological replicates.

### CRWN1-chromatin interactions mediate gene suppression

Next, we sought to explore features of gene expression associated with the dynamic CRWN1:2HA-chromatin interactions. Under both control and heat stress conditions, CRWN1:2HA showed a negative association with gene expression (Figure 4A). Among all the genes bound by CRWN1:2HA, silenced or weakly expressed genes exhibited stronger ChIP-seq signals than highly expressed ones, and the differences in ChIP-seq signal strength was located around the 5’ end of genes (Figure 4A). This observation was in line with our previous report showing that CRWN1-chromatin interactions were negatively correlated to chromatin accessibility, which was a universal feature of the 5’ end of actively expressed genes (Hu et al. 2019). We also analyzed expression changes of genes, which showed dynamic CRWN-chromatin interactions under heat stress. These genes were named “M- or H-specific” and “MH”, referring to their enrichment status with CRWN protein in mock (M) and heat stress (H) conditions. Compared to genes bound by CRWN1:2HA in both conditions (i.e., “MH”), those losing their interactions in heat (i.e., “M-specific”) tended to show elevated expression (Figure 4B). Furthermore, of the genes in the “MH” group, those annotated as upregulated DEGs (differentially expressed genes) showed attenuated interactions with CRWN1:2HA under heat stress (Figure 4C). These results reveal a correlation between dissociation of chromatin-CRWN1 and gene upregulation during heat stress response. We next asked whether CRWN1/4 target genes could be differentiated from non-target genes according to transcriptional activities. However, by comparing changes in gene expression, we did not observe any significant difference between these two groups of genes (Figure S8). Furthermore, CRWN1/4 target genes did not show a systematic shift towards up- or down-regulation in *crwn1 crwn4* mutants, implying that CRWN1/4-chromatin interactions alone are not sufficient to drive gene expression change (Figure S8).

**Figure 4.**
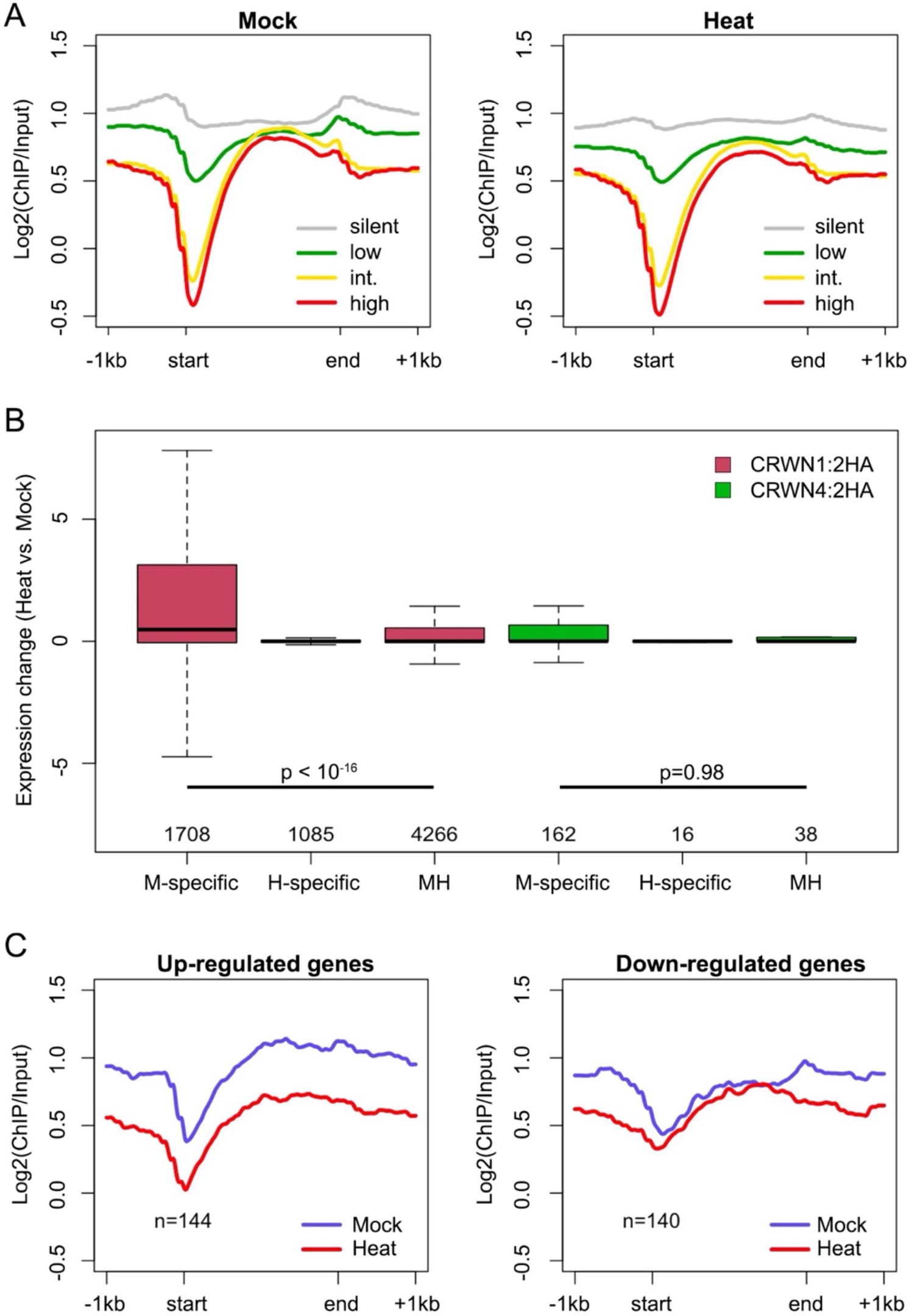
CRWN1-chromatin interactions and gene expression changes during heat stress response. (A) For CRWN1-target genes, their interaction profile with CRWN1 is shown according to gene expression levels. “int.” stands for the intermediate level. “Start” and “end” depict transcription start site and transcription termination site, respectively. (B) Changes of gene expression under heat stress. Genes enriched by CRWN only in control or heat-stressed plants are labeled as “M-specific” or “H-specific”, respectively. Genes enriched in both conditions are labeled as “MH”. The y-axis depicts differences in rpkm. p values indicate the Mann-Whitney U test results. (C) CRWN1-chromatin interactions across differentially expressed genes in the “MH” group defined in panel (B).

### CRWNs mediate chromatin organization under heat stress

The *Arabidopsis* CRWN1 and CRWN4 have been recently shown to modulate perinuclear chromatin positioning and pericentromere structure (Hu et al. 2019; Sakamoto et al. 2022). We applied the Hi-C method to examine if CRWN1 and CRWN4 play a structural role in mediating the changes of chromatin organization under heat stress (Sun et al. 2020). In total, Hi-C maps of wild-type and *crwn1 crwn4* under control and heat-stress conditions were generated (Figure S9). In line with recent studies, we observed that heat stress caused reduction of chromatin contacts within pericentromeres as well as reduction of many distal intrachromosomal contacts; additionally, we observed remarkable reduction of contact amongst KEEs (KNOT ENGAGED ELEMENTS) (Grob et al. 2014; Sun et al. 2020)(Figure S10). On the contrary, the *crwn1 crwn4* mutant Hi-C map generally showed opposite patterns, which were consist with a recent report, in which chromatin contacts within and among individual pericentromeres became stronger (Sakamoto et al. 2022)(Figure S11). In the nucleus, individual *Arabidopsis* chromosome arms fold and form two spatially insulated compartments (named compartment A/B), which have distinct gene expression and epigenetic landscape (Feng et al. 2014b; Grob et al. 2014). For chromosome arm regions, neither heat stress nor the loss of CRWNs resulted in drastic changes in the chromatin compartmentalization identity, but they both enhanced chromatin contacts between different spatial compartments (Figure 5). These changes were also clearly exhibited in heat-stressed *crwn1 crwn4* in comparison to heat-stressed wild-type plants (Figure 5). Furthermore, CRWN1 displayed preferential enrichment of chromatin binding with the B compartment, and chromatin in heat-stressed *crwn1 crwn4* showed noticeably the weakest compartmentalization, indicating that CRWNs participate in shaping genome organization under heat stress (Figure 5D and Figure 6A).

**Figure 5.**
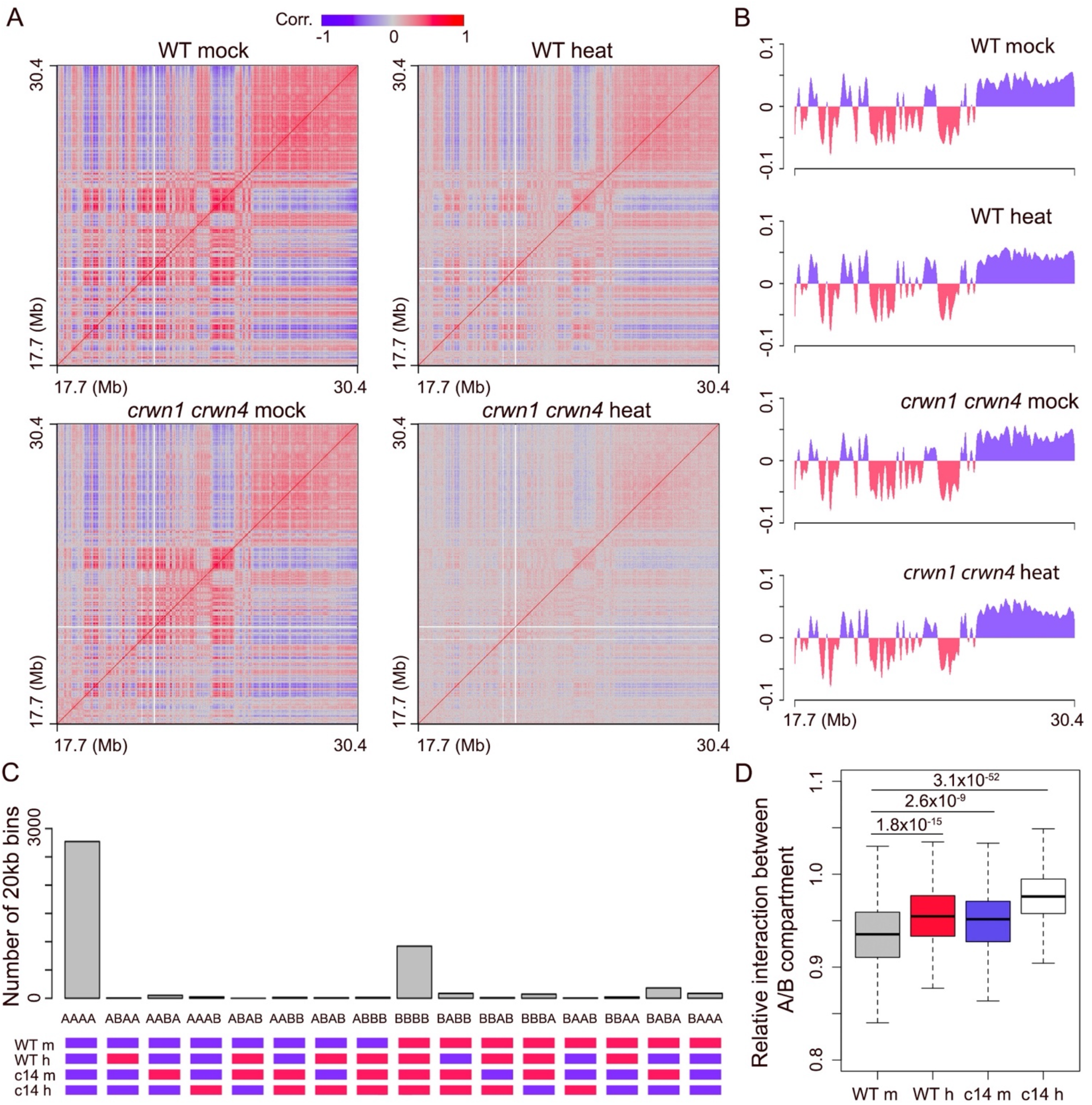
Chromatin compartments are weakened by heat and loss of CRWN1 and CRWN4. (A) Correlation matrices of Hi-C maps of chromosome 1 right arm. (B) Binary annotation of compartment A/B at chromosome 1 right arm. The y-axis shows values of the eigenvector of the first component. Regions colored in blue and red depict compartment A and B, respectively. (C) Genome-wide A/B compartment annotation among different samples. WT, wild-type; c14, *crwn1 crwn4;* m, mock; h, heat. (D) Distance normalized interactions between compartment A and B relative to the average. Sample labels are as panel C. Chromatin contacts within 1Mb are included in the calculation. p values indicate the Mann-Whitney U test results.

**Figure 6.**
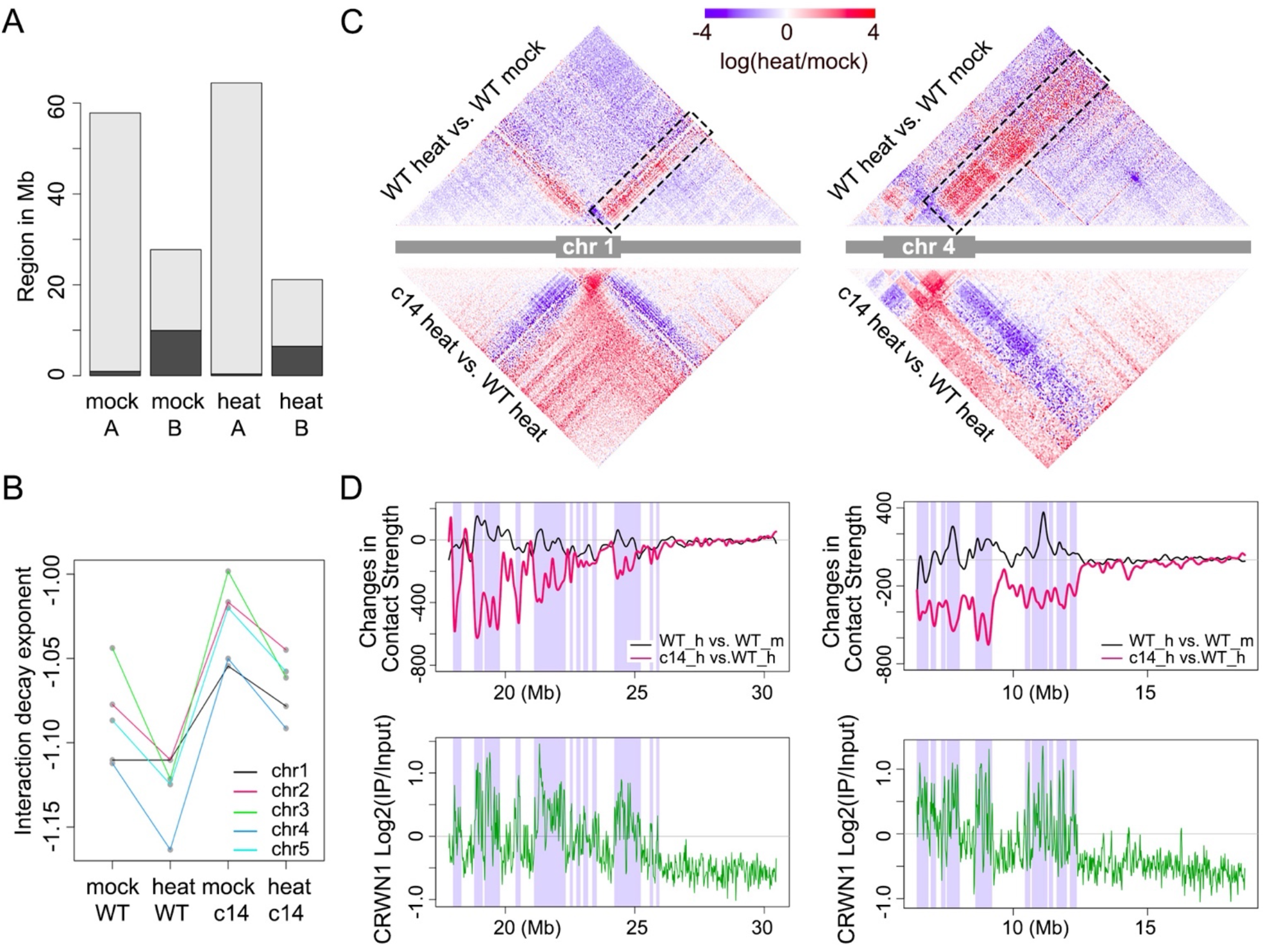
CRWN1 modulates chromatin organization under heat stress. (A) Distribution of CRWN1-enriched regions among A/B compartments in wild-type plants. (B) Interaction decay exponents of pericentromeric regions in different samples. (C) Comparison of Hi-C contacts at chromosome 1 (left) and chromosome 4 (right). Colors indicate the difference of chromatin interaction strengths, expressed as the ratio between the two selected samples. The boxes highlight areas describing altered contacts concerning interactions between pericentromeric region and chromosome arm, which are further illustrated in panel D. The pericentromeric region is depicted as a grey block in the sketch of the chromosome. (D) CRWN proteins shape chromatin organization under heat stres. Top panel, difference in contact between pericentromeric region and chromosome arm; bottom panel, CRWN1 ChIP-seq signal in heat-stressed plants. The shaded areas depict compartment B annotation in heat-stressed wild-type plants. WT, wild-type; c14, *crwn1 crwn4;* h, heat.

Another observation on pericentromeric regions further supported a structural role of CRWNs in modulating chromatin organization under heat stress. Under heat stress, pericentromeres acquired more chromatin contacts with chromosome arms along with changes in its folding conformation, which could be reflected by the IDEs (Interaction Decay Exponents) describing chromatin interaction as a function of loci distance (Figure 6B, Figure S10A)(Sun et al. 2020). Interestingly, the genomic regions showing stronger chromatin contacts with pericentromeres under heat stress tended to be associated with CRWN1 binding (Figure 6D). For *crwn1 crwn4* under heat stress, compared to wild-type plants, pericentromeres showed decreased chromatin contacts with chromosome arms, which was strongly correlated with CRWN1 ChIP-seq signals (Figure 6D). We also noticed that heat stress induced stronger CRWN1-chromatin interactions in pericentromeric regions (Figure S7A and 7C). Taken together, these results suggest that the dynamic CRWN1-chromatin interactions participate in shaping genome organization under heat stress.

## Discussion

In this work, we present data highlighting the dynamic nature of the plant nuclear lamina in response to various stress conditions such as heat, salt, and osmotic stresses (Figure 2). Since multiple nuclear lamin components interact with each other (Goto et al. 2014; Sakamoto et al. 2020), one would speculate that the assembly of the nuclear lamina relies on the presence of all these proteins. However, the periphery nuclear location of single lamin protein species turned out to be independent on other nuclear lamina components. For example, KAKU4 protein localization remains at the nuclear periphery in *crwn1 crwn4* double mutant (Goto et al. 2014). In another example, we observed that in the *crwn4* mutant background, CRWN1 proteins are still specifically located at the nuclear periphery as in wild-type plants, and they undergo disassembly in response to heat stress (data not shown). These results prompted us to speculate that the nuclear localization patterns of lamin proteins are largely independent on each other, and during stress adaptation their disassembly was triggered in parallel. To gain a better understanding on how plant nuclear lamina response to stresses, it is intriguing to further identify which part of individual lamin proteins enables their dynamic behavior.

To our surprise, the disassembly of the *Arabidopsis* nuclear lamina under heat stress does not lead to apparent changes in nuclear size and shape, albeit multiple studies have clearly demonstrated that nuclear lamin components, e.g., CRWN1, CRWN4, and KAKU4, are indispensable for proper nuclear morphology (Dittmer et al. 2007; Sakamoto and Takagi 2013; Goto et al. 2014). Although we could not verify in our experiments whether the nuclear periphery was free of nuclear lamina under stress, our observation strongly suggests that the location of tested nuclear lamin components at the nuclear periphery is not necessary in maintaining nuclear morphology. The size of animal nuclear is known to be controlled at multiple phases of a cell cycle including the nuclear envelope building stage in newly formed daughter cells (recently reviewed by (Cantwell and Dey 2021)). Our results imply that CRWNs and KAKU4 proteins play their structural roles concerning nuclear morphology before interphase; while during interphase they actively participate in mediating stress responses. In line with this notion, it has been found that both CRWN1 and CRWN4 assembles are observable at telophase when the nuclear envelope reconstruction starts (Sakamoto and Takagi 2013).

The dynamics of the *Arabidopsis* nuclear lamina suggests several future exciting research topics concerning the underlying regulatory mechanisms, spatial chromatin organization, and transcriptional regulation. First, multiple studies on CRWN1 indicate that how this plant nuclear lamin component responds to stresses in a highly variable way. CRWN1 protein start degradation after infection with virulent bacteria, which is likely mediated by the salicylic acid signaling pathway (Guo et al. 2017). On the contrary, it was shown by Sakamoto and colleagues that CRWN1 proteins remained stably localized at the nuclear periphery in roots when plants were stressed with excess copper ions (Sakamoto et al. 2020). Here, we reveal that heat stress treatment cause CRWN1 (as well as nuclear lamin components CRWN4 and KAKU4) to be redistributed in the nucleoplasm in the absence of protein degradation (Figure 2). It is unknown how CRWN1 proteins respond differently to these stress stimuli. However, based on available information on metazoan nuclear lamins (Machowska et al. 2015; Torvaldson et al. 2015), which are functionally and structurally comparable (but not sequence-wise) to the plant counterparts, we speculate that post-translational modification may confer plant nuclear lamins capability to differentiate distinct stress stimuli. Post-translational modifications of the plant nuclear lamina have not been systematically documented; nonetheless, our search over an *Arabidopsis* phosphoproteomics database (https://phosphat.uni-hohenheim.de/index.html) indicated that CRWN1, CRWN4 and KAKU4 had at between 10 to 30 experimentally confirmed phosphorylation sites (Heazlewood et al. 2008). Technically, it is feasible to profile post-translational modification patterns of a given plant nuclear lamin component under different stress conditions, which could be useful in elucidating how the nuclear lamina reacts differently to diverse stimuli.

Second, plant lamin components under various stress conditions may interact with diverse genomic regions. In this study, we focus on CRWN1 and found noticeable gain- and loss-of CRWN1-chromatin interactions during the nuclear lamina disassembly (Figure 3C), which correlated to changes in gene expression (Figure 4). Accompanied with changes in spatial distribution, the nuclear lamin proteins may switch to interact with new protein partners. It is not known if other nuclear lamina proteins, which we did not investigated in this study, interact with chromatin dependently on the status of the nuclear lamin composition. A recent proteomic approach using biotin proximity labelling showed that KAKU4 interacted with nucleosomes (presumably form chromatin) at the nuclear periphery (Tang et al. 2022). Since KAKU4 proteins detached from the nuclear envelope after perceiving stresses (Figure 2), we speculate that KAKU4 might bind to different target loci before and after nuclear lamina disassembly. In addition, whether CRWN2 and CRWN3, which are homologs of CRWN1 but located throughout the nucleoplasm (Sakamoto and Takagi 2013; Wang et al. 2013), interact directly with chromatin remains unknown. Nevertheless, we propose that this would be the case since that CRWN1 was reported to interact with all its homologs *in vivo* (Sakamoto et al. 2020).

Third, given a strong correlation between the plant nuclear periphery and suppressed gene expression (Bi et al. 2017; Hu et al. 2019), it is of great interest for us to explore to which extend changes in the nuclear lamina at the nuclear periphery are connected to transcriptional regulation in this nuclear compartment. It has been noted that a few genes showed propensity to be re-positioned to the nuclear periphery along with transcriptional activation/up-regulation, examples of which include copper-associated gene in response to copper toxicity and the CAB gene cluster (encoding chlorophyll a/b-binding proteins) in response to red/far-red light (Feng et al. 2014a; Sakamoto et al. 2020). Is such transcriptional regulation associated with changes at the nuclear periphery so that it favors transcribing these genes? Will the disassembly of plant nuclear lamina convert the nuclear periphery to a place harboring more active euchromatin? Although such a scenario has not been described in plants, case studies focusing on rod photoreceptors nuclei from nocturnal animals shows that withdrawing lamin-chromatin contacts play pivotal roles in inverting euchromatin/heterochromatin distribution patterns at the nuclear periphery (Solovei et al. 2013; Falk et al. 2019).

## Materials and Methods

### Plant Materials and growth conditions

Surface-sterilized *Arabidopsis* seeds were sowed on vertical half-strength Murashige and Skoog (1/2 MS) media plates containing 1% sucrose and 0.3% Phytagel. After being stratified at 4 °C for 3 days, the plant materials were grown in a growth chamber (MLR-352-PE from PHCbi) set at 21 °C and long day (16 h light/8h dark) condition. Mutants used in this study were *crwn1-1* (SALK_025347), *crwn4-1* (SALK_079296), and *kaku4-2* (SALK_076754), which were ordered from the Nottingham *Arabidopsis* Stock Centre.

For heat stress treatment, 11-day-old plants were transferred to another growth chamber of the same model and with identical settings, except that the temperature was set at 37 °C when lights were switched on. Plants were harvested three days later for analyses. For salt and osmotic stress treatment, 12-day-old plants were transferred to 1/2 MS plates supplemented with 150 mM Sodium Chloride and 5% Polyethylene Glycol 8000, respectively. Plants were harvested two days later for analyses.

Plant nuclei extraction from leaf tissues was performed essentially according to the steps described in our earlier work (Zhu et al. 2017). The subsequent nuclei sorting was performed with S3e Cell Sorter (Bio-Rad) according to (Wang et al. 2021). The extracted nuclei were stained with 0.5 μM DAPI to reveal their ploidy levels; if not otherwise specified, only 2C nuclei were collected for downstream experiments.

### Plasmid Construction

For the *CRWN4:2HA* construct, a tandem HA tag (2HA) was inserted after the 850th amino residue of the CRWN4 protein, generating a *CRWN4:2HA* fusing construct that fully rescued *crwn4* loss-of-function mutant phenotypes. The two fragments of this *CRWN4:2HA* construct were amplified with primers: 5’-ACTAATCTTTTCTACTAGCTTAAC -3’ in combination with 5’-AGGGTATCCAGCATAATCTGGTACGTCGTATGGGTATCCAGTACATCGTTTTATCCATGA -3’; and 5’-GATTATGCTGGATACCCTTACGACGTACCAGATTACGCTAATCTGATTTTCAAGACTTCTCCA -3’ in combination with 5’-GCTACGAGCTACTTCGATGATAC -3’, respectively. These two fragments, which collectively cover the genomic fragment of *CRWN4* locus and its 2 kb promoter regions, were assembled with overlapping PCR and subsequently amplified with primers 5’-ACTAATCTTTTCTACTAGCTTAAC -3’ and 5’-GCTACGAGCTACTTCGATGATAC -3’. The PCR product was cloned into the pFK206 vector (Karlsson et al. 2015).

For the *KAKU4:GFP* construct, the genomic region containing 2kb upstream of *KAKU4* and its coding sequence was amplified with oligos 5’-GCATAGAACGAGGAATACAGG -3’ and 5’-CTGCCTCCTGCAGCTCCGGATTTGGCCCGTCCTTTGCCTC -3’, and the GFP cDNA was amplified with oligos 5’-TCCGGAGCTGCAGGAGGCAGCGCGGCCGCTGTGAGCAAGGG-3’ and 5’-TTATCCGGACTTGTACAGCTCG-3’. These two PCR fragments were purified and cloned into the pFK206 vector with a gibson assembly reaction, by which the GFP sequence was appended to the C-terminus of KAKU4.

### Fluorescence in Situ Hybridization (FISH) and Immunohistostaining

BAC (Bacterial Artificial Chromosome) probes were generated by the nick translation DNA labeling system (Roche, Cat. No. 11745808910). The green and red probes described in this study were labeled by Digoxigenin (DIG) and 2,4-Dinitrophenyl (DNP), respectively. The labeled BACs were pooled in a hybridization mix (50% formamide, 10% dextran sulfate, 2× SSC, 50 mM sodium phosphate (pH 7.0)]. The working concentration of each labeled BAC was 1 ng/*μ*l. Individual BACs were listed in Supplemental Table 1.

FISH experiments were performed according to (Montgomery et al. 2022). Briefly, around 5000 nuclei in 20 *μl* PBS buffer were incubated at 65°C for 30 min. The nuclei were subsequently mixed with 10 *μ*l of 0.1 mg/ml RNase A and spread into a circle on a glass slide drawn by an ImmEdge™ pen. After incubating the slide at 37 °C in a hybridizer (ThermoBrite, model 07J91-020) for 1 hour, the slide were treated for 1min each by dipping up and down in an ethanol series (30 %, 60 %, 80 %, 90 %, 95 %, 100 % EtOH) to dehydrate. For immunohistostaining, an antigen retrieval step was performed by incubating the slides in boiling solution (10 mM sodium citrate at pH 6.0) for 12 min in a microwave at 700 W. After antigen retrieval, slides were post-fixated in 4 % formaldehyde solution for 10 min, followed by a series of ethanol dehydration treatment, and then air-dried. The treated slides were proceeded to FISH or immunohistostaining experiments. For FISH experiment, the probe hybridization, slides washing, and signal detection steps were performed according to (Bi et al. 2017) with minor changes. DIG-labeled probes were detected with 1:10 diluted digoxigenin Alexa Fluor 488-conjugated mouse antibody (Biotechne, Catalog no. IC7520G), and DNP-labeled probes were detected with 1:500 diluted DNP rabbit antibody (ThermoFisher, Catalog no. 04-8300) and 1:150 diluted Anti-rabbit Alexa Fluor 546-conjugated goat antibody (ThermoFisher, Catalog no. A-11035). For immunohistostaining, the HA-tagged protein of interest was detected with 1:500 diluted HA Tag Alexa Fluor 647 conjugated mouse antibody (ThermoFisher, Catalog no. 26183-A647). After antibody incubation, the slides were washed with 4x SSC 0.2 % tween 20 in a foil-wrapped jar at room temperature 5 minutes for three times. Finally, slides were mounted with 5 *μ*l SlowFade™ Diamond Antifade Mountant (Invitrogen, Catalog no. S36964).

### Microscopy and Image Processing

Confocal images were captured with a Zeiss LSM 700 system. For FISH experiments, a single image was taken from the central focal plane of individual nuclei. Image analyses were performed with ImageJ (Schneider et al. 2012). The distance between a FISH signal spot and the nuclear periphery was recorded as the distance between its estimated barycenter and the edge of DAPI staining. We noticed that a fraction of the observed nuclei showed split or scattered FISH signal patterns, making it difficult to estimate the distance between the probed genomic region to the nuclear envelope (Figure S1). These nuclei were excluded from analysis. Similarly, an image of the central focal plane of a nucleus was taken to analyze the distribution of the protein of interest. Images of chromosome painting were acquired with a Zeiss LSM 880 system. Quantification of nuclear size was performed with images of DAPI-stained nuclei taken with an Olympus IX83 fluorescence microscope. The Olympus cellSens software was used to measure the area of nuclei in the images, which was used as an approximation to nuclear size.

### Protein extraction and western blot

Protein extraction from aerial tissues was performed with sample homogenization using protein extraction buffer (50 mM Tris-HCl, 150 mM NaCl, 0.1% Tween 20) supplemented with 1% β-Mercaptoethanol (β-ME), 0.1 M Phenylmethylsulfonylfluoride, and Protease Inhibitor (Protease Inhibitor Cocktail Tablets; Roche). The homogenate was centrifuged for 10 minutes at 4° C at 12000 rpm. The supernatant was recovered and analyzed by western blot. The following antibodies were used to detect protein of interest: for HA-tagged proteins, anti-HA-HRP (sc-7392, Santa Cruz Biotechnology); for GFP-tagged proteins, anti-GFP (anti-GFP (ab290, Abcam) followed by anti-rabbit HRP conjugate (A6154, Sigma-Aldrich). After chemiluminescence detection, membranes were stained with coomassie blue.

### Chromatin immunoprecipitation and library sequencing

*Arabidopsis* shoots were fixed under vacuum for 30 min with 1% formaldehyde in MC buffer (10 mM potassium phosphate, pH 7.0, 50 mM NaCl, 0.1 M sucrose) at room temperature. Fixation was terminated by replacing the solution with 0.15 M glycine in MC buffer under vacuum for 10 min at room temperature. Approximately 1 g fixed tissue was homogenized and resuspended in nuclei isolation buffer (20 mM Hepes, pH 8.0, 250 mM sucrose, 1 mM MgCl_2_, 5 mM KCl, 40% glycerol, 0.25% Triton X-100, 0.1 mM PMSF, 0.1% 2-mercaptoethanol) and filtered with double-layered miracloth (Millipore). Isolated nuclei were resuspended in 0.5 mL sonication buffer (10 mM potassium phosphate, pH 7.0, 0.1 mM NaCl, 0.5% sarkosyl, 10 mM EDTA), and chromatin was sheared by sonication with a QSONICA sonicator Q800R3 to achieve average fragment size around 400 bp. Next, 50 μL 10% Triton X-100 was mixed with the sonicated sample, and 25 μl of the mixture was saved as input sample. The rest of the sheared chromatin was mixed with an equal volume of IP buffer (50 mM Hepes, pH 7.5, 150 mM NaCl, 5mM MgCl2, 10 μM ZnSO4, 1% Triton X-100, 0.05% SDS) and incubated with PierceTM Anti-HA magnetic beads (Thermo Fisher) at 4°C for 2h. The beads were washed at 4°C as follows: 2× with IP buffer, 1× with IP buffer having 500 mM NaCl, and 1× with LiCl buffer (0.25 M LiCl, 1% NP-40, 1% deoxycholate, 1mM EDTA, 10 mM Tris pH 8.0) for 3 min each. After a brief wash with TE buffer (10 mM Tris pH 8.0, 1mM EDTA), the beads were resuspended in 200 μL elution buffer (50 mM Tris, pH 8.0, 200 mM NaCl, 1% SDS, 10 mM EDTA) at 65°C for 6 h, followed by Proteinase K treatment at 45°C for 1 h. DNA was purified with MinElute PCR purification kit (QIAGEN), and then used for qPCR or converted into sequencing libraries following the NEBNext^®^ Ultra™ II DNA Library Prep Kit (NEB). For ChIP-qPCR, the relative enrichment of tested loci was normalized to TUB2. Oligos used for ChIP-qPCR are listed in Supplemental Table 2.

### RNA-seq Library Preparation and analysis

RNA sequencing (RNA-Seq) was performed with two biological replicates per sample. Total RNA was extracted from aerial parts of seedlings using a RNeasy Plant Mini Kit (Qiagen). RNA-seq library preparation was performed as described (Wang et al. 2021).

RNA-seq sequencing reads were aligned against the Araport11 annotation using TopHat 2 (v2.1.1) with default parameters, and were further assigned to genes using the R package GenomicAlignments (Kim et al. 2013; Lawrence et al. 2013; Cheng et al. 2017). Differentially expressed genes were identified with the R package DESeq2 (Love et al. 2014). We used criteria of false discovery rate smaller than 0.01 and log2 expression fold change more than 1.6 to call up- and down-regulated genes. Details of the reads count table, gene expression measurement (in reads per kilobase per million mapped reads), and differentially expressed genes can be found in Supplemental Table 3.

### *In situ* Hi-C

In situ Hi-C library preparation was followed essentially as described (Karaaslan et al. 2020). In total, two Hi-C library replicates for each sample were made, and for each replicate around 0.5 g of fixed sample were homogenized for nuclei isolation. Nuclei were resuspended with 150 μl 0.5% SDS and split into three tubes. After penetration at 62°C for 5 min, SDS was quenched by adding 145 μl water and 25 μl 10% Triton X-100, and incubated at 37°C for 15 min. Subsequently, chromatin was digested overnight at 37°C with 50 U Dpn II (NEB) in each tube. The next day, Dpn II was inactivated by incubating at 62°C for 20 min. Then, sticky ends were filled in by adding 1 μl of 10 mM dTTP, 1 μl of 10 mM dATP, 1 μl of 10 mM dGTP, 10 μl of 1 mM biotin-14-dCTP, 29 μl water and 40 U Klenow (Thermo Fisher), and incubated at 37 °C for 2 h. After adding 663 μl water, 120 μl 10x blunt-end ligation buffer (300 mM Tris-HCl, 100 mM MgCl2, 100 mM DTT, 1 mM ATP, pH 7.8) and 40 U T4 DNA ligase (Thermo Fisher), proximity ligation was carried out at room temperature for 4 hours. Then, three tubes of ligation products were centrifuged, and nuclei pellet was resuspended and combined with 650 μl SDS buffer (50 mM Tris-HCl, 1% SDS, 10 mM EDTA, pH 8.0). After treatment with 10 μl proteinase K (Thermo Fisher) at 55 °C for 30 min, decrosslinking was performed by adding 30 *μ*l 5 M NaCl, and incubated at 65 °C overnight.

DNA was recovered and subsequently treated with RNase A at 37 °C for 30 min. After purification, 3~5 μg Hi-C DNA was topped to 130 μl with TE buffer (10 mM Tris-HCl, 1 mM EDTA, pH 8.0), and sheared with Q800R3 (QSONICA) sonicator by using the following setting: 25% amplitude, 15s ON, 15s OFF, pulse-on time for 4.5 min, to achieve fragment size shorter than 500 bp. Sonicated DNA was purified with Ampure beads to recover fragments longer than 300 bp. Then, with a 50 μl reaction volume, the DNA was mixed with 0.5 μl 10 mM dTTP, 0.5 μl 10 mM dATP, and 5 U T4 DNA polymerase and incubated at 20 °C for 30 min to remove biotin from unligated DNA ends. After that, the DNA was purified with Ampure beads, continued with end-repair and adaptor ligation by following the NEBNext^®^ Ultra™ II DNA Library Prep Kit (NEB). Ligated DNA was affinity-purified with Dynabeads MyOne Streptavidin C1 beads (Invitrogen) as described (Liu et al. 2016), and further amplified with 12 PCR cycles. The libraries were sequenced on an Illumina Novaseq instrument with 2 x 150 bp reads.

Reads mapping to the TAIR10 genome, removal of PCR duplicates, and reads filtering were performed as described (Liu et al. 2016). Hi-C reads of each sample are summarized in Supplemental Table 4. Hi-C map normalization was performed by using an iterative matrix correction function in the “HiTC” package in R programm (Servant et al. 2012). For all Hi-C maps, the iterative normalization process was stopped when the eps value, which reflected how similar the matrices in two consecutive correction steps were, dropped below 1 × 10^-4^. The bin size setting for genome-wide Hi-C maps was 20 kb. Besides, the filtered Hi-C reads were used to create *hic* files with the juicer tool (Durand et al. 2016) for interactive Hi-C map inspection. Genomic coordinates of individual pericentromeric regions for computing IDE (interaction decay exponent) were defined as follows: Chr 1: 11.5-17.7 Mb; Chr 2: 1.1-7.2 Mb; Chr 3: 10.3-17.3 Mb; Chr 4: 1.5-6.3 Mb; and Chr 5: 9.0-16.0 Mb (Stroud et al. 2013).

## Data availability

Short read data of in situ Hi-C, ChIP-seq, and RNA-seq are publicly available at NCBI Sequence Read Archive under accession number PRJNA870030.

Large datasets, such as Hi-C matrices and ChIP-seq track files in 100 bp bin size are available in the figshare repository, which are accessible with the following link: https://figshare.com/s/8dc4d77ca579b73bbbe4 with Digital Object Identifier (DOI) 10.6084/m9.figshare.21370560.

## Acknowledgements

We thank Marie-Edith Chabouté for providing us *pSUN1:SUN1-GFP* seeds. We thank computing support by the High Performance and Cloud Computing Group at the Zentrum für Datenverarbeitung of the University of Tübingen, the state of Baden-Württemberg through bwHPC and the German Research Foundation (DFG) through grant no. INST 37/935-1 FUGG. We thank Sandra Richter and Natalie Krieger from the microscopy department at the ZMBP, University of Tübingen for their assistance in LSM880 usage. We thank Kerstin Feistel from the Department of Zoology, University of Hohenheim for her assistance in image acquisition with LSM700.

## References

Barneche F, Malapeira J, Mas P. 2014. The impact of chromatin dynamics on plant light responses and circadian clock function. J Exp Bot 65: 2895–2913.

Bhadouriya SL, Mehrotra S, Basantani MK, Loake GJ, Mehrotra R. 2020. Role of Chromatin Architecture in Plant Stress Responses: An Update. Front Plant Sci 11: 603380.

Bi X, Cheng YJ, Hu B, Ma X, Wu R, Wang JW, Liu C. 2017. Nonrandom domain organization of the Arabidopsis genome at the nuclear periphery. Genome Res 27: 1162–1173.

Bourbousse C, Mestiri I, Zabulon G, Bourge M, Formiggini F, Koini MA, Brown SC, Fransz P, Bowler C, Barneche F. 2015. Light signaling controls nuclear architecture reorganization during seedling establishment. Proc Natl Acad Sci U S A 112: E2836–2844.

Cantwell H, Dey G. 2021. Nuclear size and shape control. Semin Cell Dev Biol doi:10.1016/j.semcdb.2021.10.013.

Cheng CY, Krishnakumar V, Chan AP, Thibaud-Nissen F, Schobel S, Town CD. 2017. Araport11: a complete reannotation of the Arabidopsis thaliana reference genome. Plant J 89: 789–804.

Choi J, Strickler SR, Richards EJ. 2019. Loss of CRWN Nuclear Proteins Induces Cell Death and Salicylic Acid Defense Signaling. Plant Physiol 179: 1315–1329.

Ciska M, Hikida R, Masuda K, Moreno Diaz de la Espina S. 2019. Evolutionary history and structure of nuclear matrix constituent proteins, the plant analogues of lamins. J Exp Bot 70: 2651–2664.

Ciska M, Masuda K, Moreno Diaz de la Espina S. 2013. Lamin-like analogues in plants: the characterization of NMCP1 in Allium cepa. J Exp Bot 64: 1553–1564.

Ciska M, Masuda K, Moreno Diaz de la Espina S. 2018. Characterization of the lamin analogue NMCP2 in the monocot Allium cepa. Chromosoma 127: 103–113.

Dittmer TA, Stacey NJ, Sugimoto-Shirasu K, Richards EJ. 2007. LITTLE NUCLEI genes affecting nuclear morphology in Arabidopsis thaliana. Plant Cell 19: 2793–2803.

Durand NC, Shamim MS, Machol I, Rao SS, Huntley MH, Lander ES, Aiden EL. 2016. Juicer Provides a One-Click System for Analyzing Loop-Resolution Hi-C Experiments. Cell Syst 3: 95–98.

Falk M, Feodorova Y, Naumova N, Imakaev M, Lajoie BR, Leonhardt H, Joffe B, Dekker J, Fudenberg G,Solovei I et al. 2019. Heterochromatin drives compartmentalization of inverted and conventional nuclei. Nature 570: 395–399.

Feng CM, Qiu Y, Van Buskirk EK, Yang EJ, Chen M. 2014a. Light-regulated gene repositioning in Arabidopsis. Nat Commun 5: 3027.

Feng S, Cokus SJ, Schubert V, Zhai J, Pellegrini M, Jacobsen SE. 2014b. Genome-wide Hi-C analyses in wild-type and mutants reveal high-resolution chromatin interactions in Arabidopsis. Mol Cell 55: 694–707.

Goto C, Tamura K, Fukao Y, Shimada T, Hara-Nishimura I. 2014. The Novel Nuclear Envelope Protein KAKU4 Modulates Nuclear Morphology in Arabidopsis. Plant Cell 26: 2143–2155.

Grob S, Schmid MW, Grossniklaus U. 2014. Hi-C analysis in Arabidopsis identifies the KNOT, a structure with similarities to the flamenco locus of Drosophila. Mol Cell 55: 678–693.

Groves NR, Biel A, Moser M, Mendes T, Amstutz K, Meier I. 2020. Recent advances in understanding the biological roles of the plant nuclear envelope. Nucleus 11: 330–346.

Gruenbaum Y, Margalit A, Goldman RD, Shumaker DK, Wilson KL. 2005. The nuclear lamina comes of age. Nat Rev Mol Cell Biol 6: 21–31.

Guo T, Mao X, Zhang H, Zhang Y, Fu M, Sun Z, Kuai P, Lou Y, Fang Y. 2017. Lamin-like Proteins Negatively Regulate Plant Immunity through NAC WITH TRANSMEMBRANE MOTIF1-LIKE9 and NONEXPRESSOR OF PR GENES1 in Arabidopsis thaliana. Mol Plant 10: 1334–1348.

Heazlewood JL, Durek P, Hummel J, Selbig J, Weckwerth W, Walther D, Schulze WX. 2008. PhosPhAt: a database of phosphorylation sites in Arabidopsis thaliana and a plant-specific phosphorylation site predictor. Nucleic Acids Res 36: D1015–1021.

Hu B, Wang N, Bi X, Karaaslan ES, Weber AL, Zhu W, Berendzen KW, Liu C. 2019. Plant lamin-like proteins mediate chromatin tethering at the nuclear periphery. Genome Biol 20: 87.

Huang A, Tang Y, Shi X, Jia M, Zhu J, Yan X, Chen H, Gu Y. 2020. Proximity labeling proteomics reveals critical regulators for inner nuclear membrane protein degradation in plants. Nat Commun 11: 3284.

Karaaslan ES, Wang N, Faiss N, Liang Y, Montgomery SA, Laubinger S, Berendzen KW, Berger F, Breuninger H, Liu C. 2020. Marchantia TCP transcription factor activity correlates with three-dimensional chromatin structure. Nat Plants 6: 1250–1261.

Karlsson P, Christie MD, Seymour DK, Wang H, Wang X, Hagmann J, Kulcheski F, Manavella PA. 2015. KH domain protein RCF3 is a tissue-biased regulator of the plant miRNA biogenesis cofactor HYL1. Proc Natl Acad Sci U S A 112: 14096–14101.

Kim D, Pertea G, Trapnell C, Pimentel H, Kelley R, Salzberg SL. 2013. TopHat2: accurate alignment of transcriptomes in the presence of insertions, deletions and gene fusions. Genome Biol 14: R36.

Kimura Y, Kuroda C, Masuda K. 2010. Differential nuclear envelope assembly at the end of mitosis in suspension-cultured Apium graveolens cells. Chromosoma 119: 195–204.

Lawrence M, Huber W, Pages H, Aboyoun P, Carlson M, Gentleman R, Morgan MT, Carey VJ. 2013. Software for computing and annotating genomic ranges. PLoS Comput Biol 9: e1003118.

Liu C, Wang C, Wang G, Becker C, Zaidem M, Weigel D. 2016. Genome-wide analysis of chromatin packing in Arabidopsis thaliana at single-gene resolution. Genome Res 26: 1057–1068.

Love MI, Huber W, Anders S. 2014. Moderated estimation of fold change and dispersion for RNA-seq data with DESeq2. Genome Biol 15: 550.

Machowska M, Piekarowicz K, Rzepecki R. 2015. Regulation of lamin properties and functions: does phosphorylation do it all? Open Biol 5.

Masuda K, Hikida R, Fujino K. 2021. The plant nuclear lamina proteins NMCP1 and NMCP2 form a filamentous network with lateral filament associations. J Exp Bot 72: 6190–6204.

Mathieu O, Jasencakova Z, Vaillant I, Gendrel AV, Colot V, Schubert I, Tourmente S. 2003. Changes in 5S rDNA chromatin organization and transcription during heterochromatin establishment in Arabidopsis. Plant Cell 15: 2929–2939.

McKenna JF, Gumber HK, Turpin ZM, Jalovec AM, Kartick AC, Graumann K, Bass HW. 2021. Maize (Zea mays L.) Nucleoskeletal Proteins Regulate Nuclear Envelope Remodeling and Function in Stomatal Complex Development and Pollen Viability. Front Plant Sci 12: 645218.

Montgomery SA, Hisanaga T, Wang N, Axelsson E, Akimcheva S, Sramek M, Liu C, Berger F. 2022. Polycomb-mediated repression of paternal chromosomes maintains haploid dosage in diploid embryos of Marchantia. Elife 11.

Ostlund C, Folker ES, Choi JC, Gomes ER, Gundersen GG, Worman HJ. 2009. Dynamics and molecular interactions of linker of nucleoskeleton and cytoskeleton (LINC) complex proteins. J Cell Sci 122: 4099–4108.

Pavet V, Quintero C, Cecchini NM, Rosa AL, Alvarez ME. 2006. Arabidopsis displays centromeric DNA hypomethylation and cytological alterations of heterochromatin upon attack by pseudomonas syringae. Mol Plant Microbe Interact 19: 577–587.

Pawar V, Poulet A, Detourne G, Tatout C, Vanrobays E, Evans DE, Graumann K. 2016. A novel family of plant nuclear envelope-associated proteins. J Exp Bot 67: 5699–5710.

Pecinka A, Dinh HQ, Baubec T, Rosa M, Lettner N, Mittelsten Scheid O. 2010. Epigenetic regulation of repetitive elements is attenuated by prolonged heat stress in Arabidopsis. Plant Cell 22: 3118–3129.

Sakamoto T, Sakamoto Y, Grob S, Slane D, Yamashita T, Ito N, Oko Y, Sugiyama T, Higaki T,Hasezawa S et al. 2022. Two-step regulation of centromere distribution by condensin II and the nuclear envelope proteins. Nat Plants doi:10.1038/s41477-022-01200-3.

Sakamoto Y, Sato M, Sato Y, Harada A, Suzuki T, Goto C, Tamura K, Toyooka K, Kimura H,Ohkawa Y et al. 2020. Subnuclear gene positioning through lamina association affects copper tolerance. Nat Commun 11: 5914.

Sakamoto Y, Takagi S. 2013. LITTLE NUCLEI 1 and 4 regulate nuclear morphology in Arabidopsis thaliana. Plant Cell Physiol 54: 622–633.

Schneider CA, Rasband WS, Eliceiri KW. 2012. NIH Image to ImageJ: 25 years of image analysis. Nat Methods 9: 671–675.

Servant N, Lajoie BR, Nora EP, Giorgetti L, Chen CJ, Heard E, Dekker J, Barillot E. 2012. HiTC: exploration of high-throughput ‘C’ experiments. Bioinformatics 28: 2843–2844.

Solovei I, Wang AS, Thanisch K, Schmidt CS, Krebs S, Zwerger M, Cohen TV, Devys D, Foisner R,Peichl L et al. 2013. LBR and lamin A/C sequentially tether peripheral heterochromatin and inversely regulate differentiation. Cell 152: 584–598.

Stroud H, Greenberg MV, Feng S, Bernatavichute YV, Jacobsen SE. 2013. Comprehensive analysis of silencing mutants reveals complex regulation of the Arabidopsis methylome. Cell 152: 352–364.

Sun L, Jing Y, Liu X, Li Q, Xue Z, Cheng Z, Wang D, He H, Qian W. 2020. Heat stress-induced transposon activation correlates with 3D chromatin organization rearrangement in Arabidopsis. Nat Commun 11: 1886.

Tamura K, Fukao Y, Iwamoto M, Haraguchi T, Hara-Nishimura I. 2010. Identification and characterization of nuclear pore complex components in Arabidopsis thaliana. Plant Cell 22: 4084–4097.

Tang Y, Dong Q, Wang T, Gong L, Gu Y. 2022. PNET2 is a component of the plant nuclear lamina and is required for proper genome organization and activity. Dev Cell 57: 19–31 e16.

Tessadori F, Chupeau MC, Chupeau Y, Knip M, Germann S, van Driel R, Fransz P, Gaudin V. 2007a. Large-scale dissociation and sequential reassembly of pericentric heterochromatin in dedifferentiated Arabidopsis cells. J Cell Sci 120: 1200–1208.

Tessadori F, Schulkes RK, van Driel R, Fransz P. 2007b. Light-regulated large-scale reorganization of chromatin during the floral transition in Arabidopsis. Plant J 50: 848–857.

Torvaldson E, Kochin V, Eriksson JE. 2015. Phosphorylation of lamins determine their structural properties and signaling functions. Nucleus 6: 166–171.

van Zanten M, Koini MA, Geyer R, Liu Y, Brambilla V, Bartels D, Koornneef M, Fransz P, Soppe WJ. 2011. Seed maturation in Arabidopsis thaliana is characterized by nuclear size reduction and increased chromatin condensation. Proc Natl Acad Sci U S A 108: 20219–20224.

Wang H, Dittmer TA, Richards EJ. 2013. Arabidopsis CROWDED NUCLEI (CRWN) proteins are required for nuclear size control and heterochromatin organization. BMC Plant Biol 13: 200.

Wang N, Karaaslan ES, Faiss N, Berendzen KW, Liu C. 2021. Characterization of a Plant Nuclear Matrix Constituent Protein in Liverwort. Front Plant Sci 12: 670306.

Yang J, Chang Y, Qin Y, Chen D, Zhu T, Peng K, Wang H, Tang N, Li X,Wang Y et al. 2020. A lamin-like protein OsNMCP1 regulates drought resistance and root growth through chromatin accessibility modulation by interacting with a chromatin remodeller OsSWI3C in rice. New Phytol 227: 65–83.

Zhu W, Hu B, Becker C, Dogan ES, Berendzen KW, Weigel D, Liu C. 2017. Altered chromatin compaction and histone methylation drive non-additive gene expression in an interspecific Arabidopsis hybrid. Genome Biol 18: 157.

