## supplemental files for "The plant nuclear lamina disassembles to regulate genome folding in stress conditions"

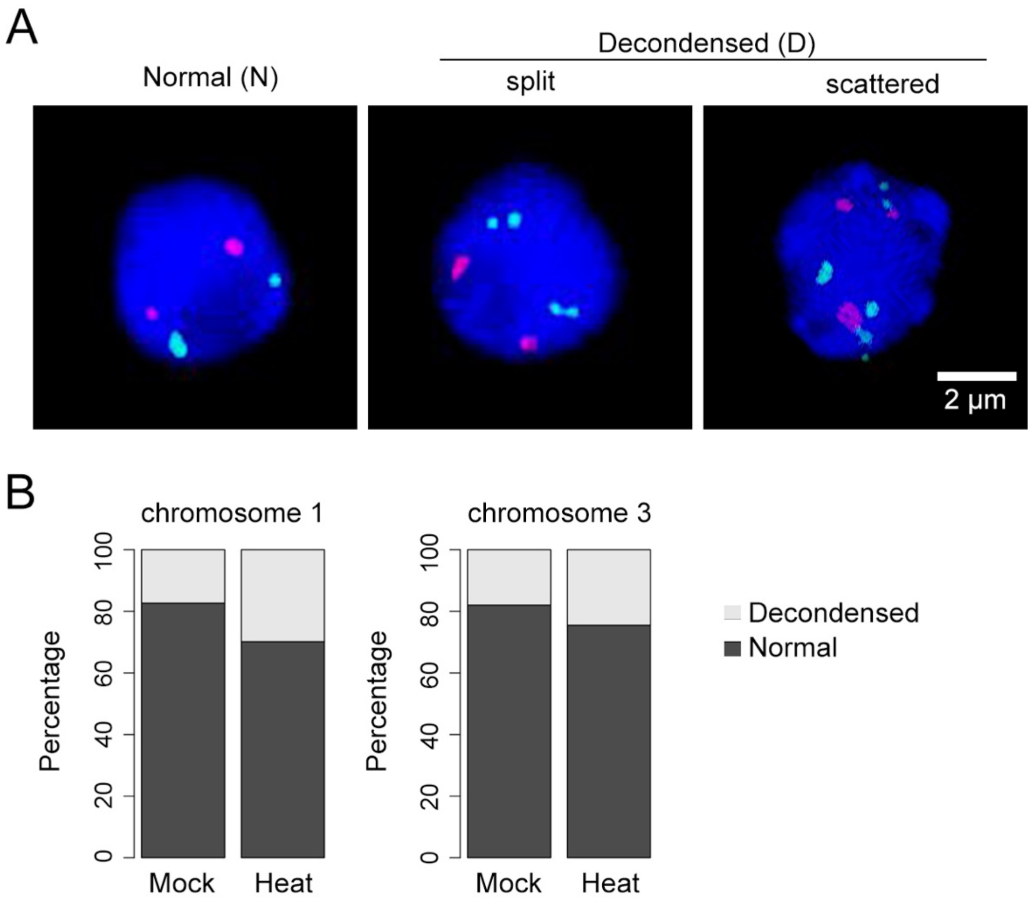


**Supplemental Figure 1. Categorization of FISH patterns in nuclei isolated from mock- and heat stress-treated plants.**
(A) Representative images of LAD (green) and non-LAD (red) probes. FISH signals indicating decondensed genomic regions were not included in distance measurement, which is shown in Figure 1C. (B) Percentage of individual categories.

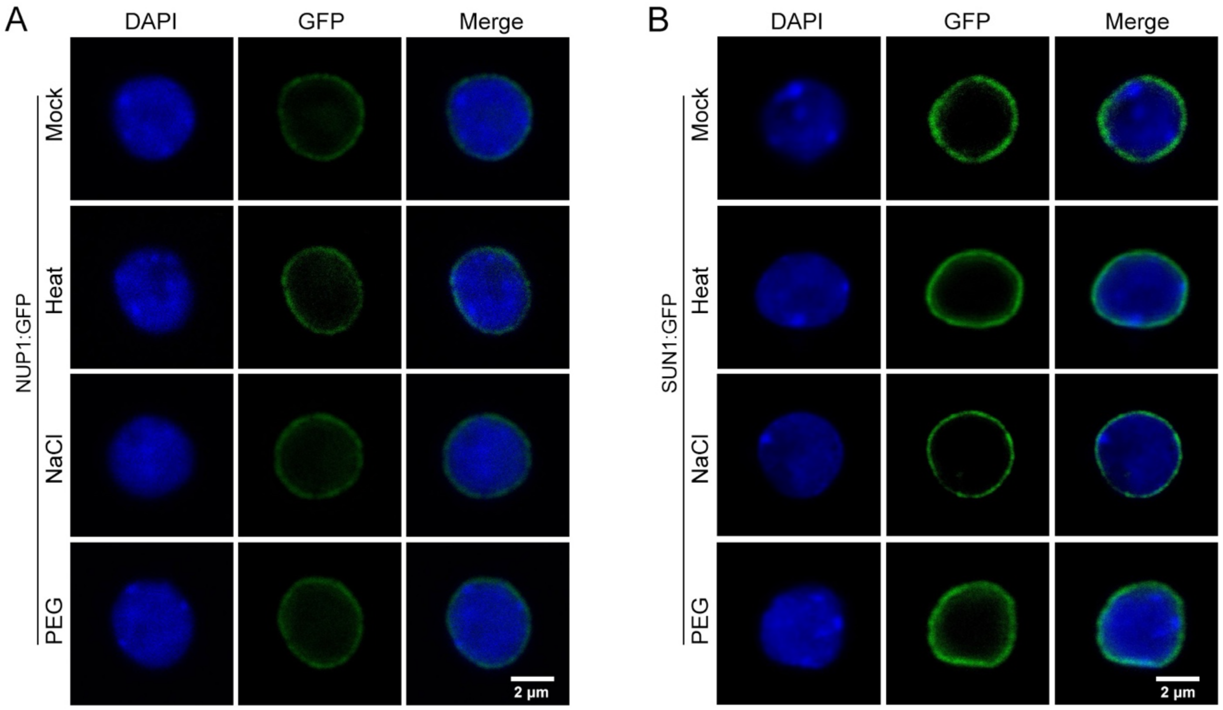


**Supplemental Figure 2. Localization of selected nuclear peripheral proteins in various growth conditions.**

(A, B) Representative confocal images of NUP1:GFP (A) and SUN1:GFP (B) in nuclei isolated from plants treated with different abiotic stresses, which are described in Figure 2.

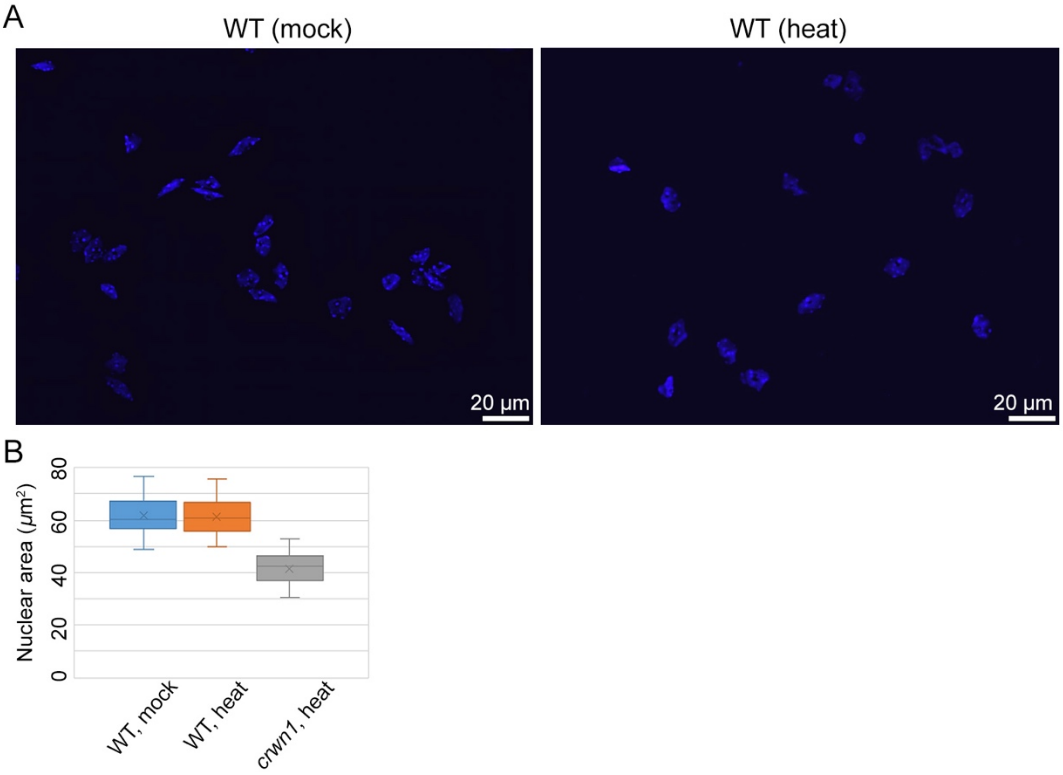


**Supplemental Figure 3. Heat stress does not influence nuclear size.**
(A) Comparison of 8C nuclei isolated from heat-stressed and control plant leaves. (B) Comparison of nuclear size, which is approximated as the area occupied by a nucleus in the microscope image. Mann-Whitney U test results comparing wild-type nuclei in mock and heat samples indicated a *p*-value larger than 0.05.

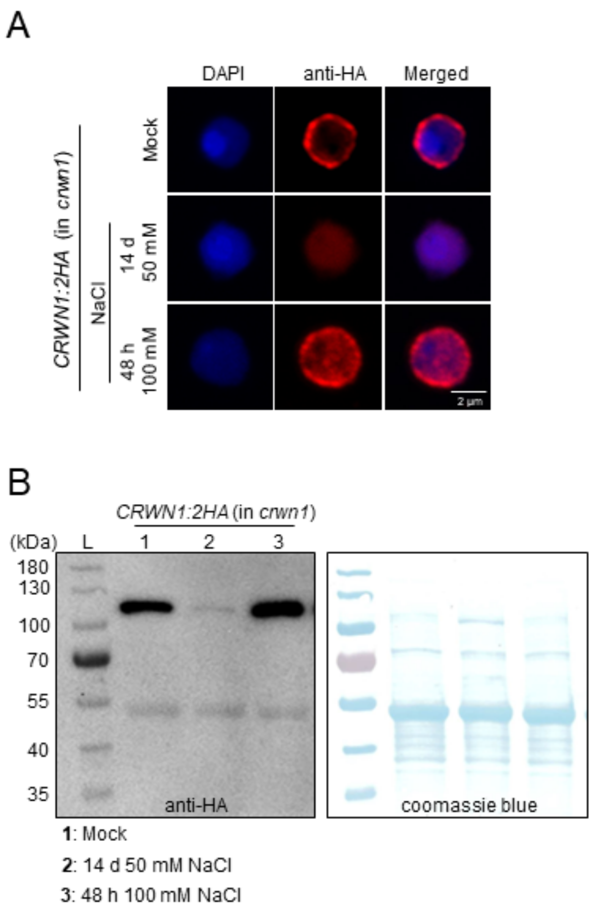


**Supplemental Figure 4. Distinct responses of CRWN1 proteins to different salt stress conditions.**
(A) Immunohistostaining of CRWN1:2HA in nuclei. (B) Immunoblots of CRWN1:2HA proteins extracted from leaves. L, protein ladder.

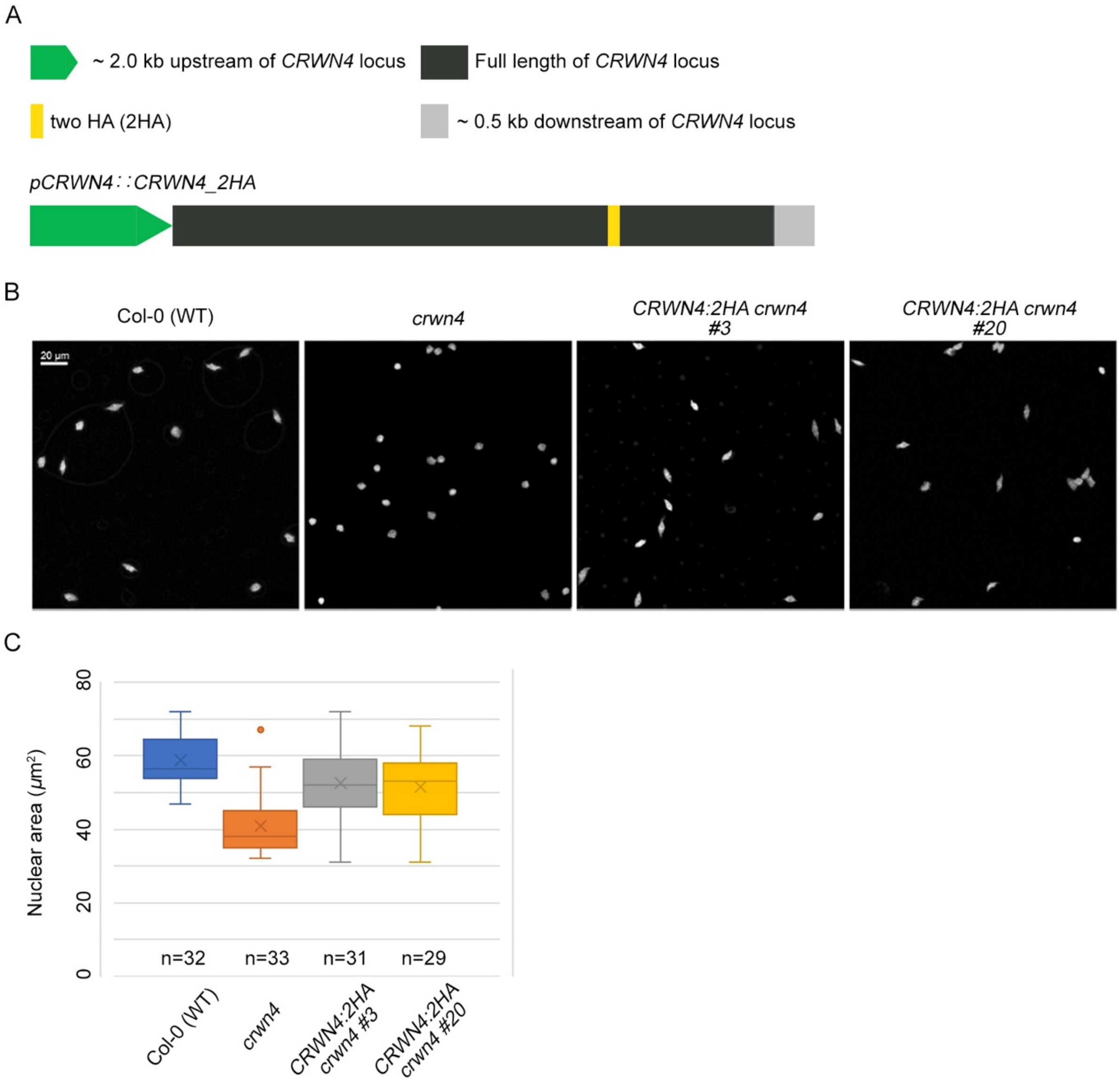


**Supplemental Figure 5. Functional complementation of *crwn4* phenotypes by the *CRWN4:2HA* transgene.**
(A) A sketch showing the *pCRWN4:CRWN4:2HA* transgene. (B) Comparison of nuclear morphology of 8C nuclei of wild-type, *crwn4*, and two lines of *pCRWN4:CRWN4:2HA* plants. (C) Boxplots of nuclear sizes of 8C nuclei isolated from different plants.

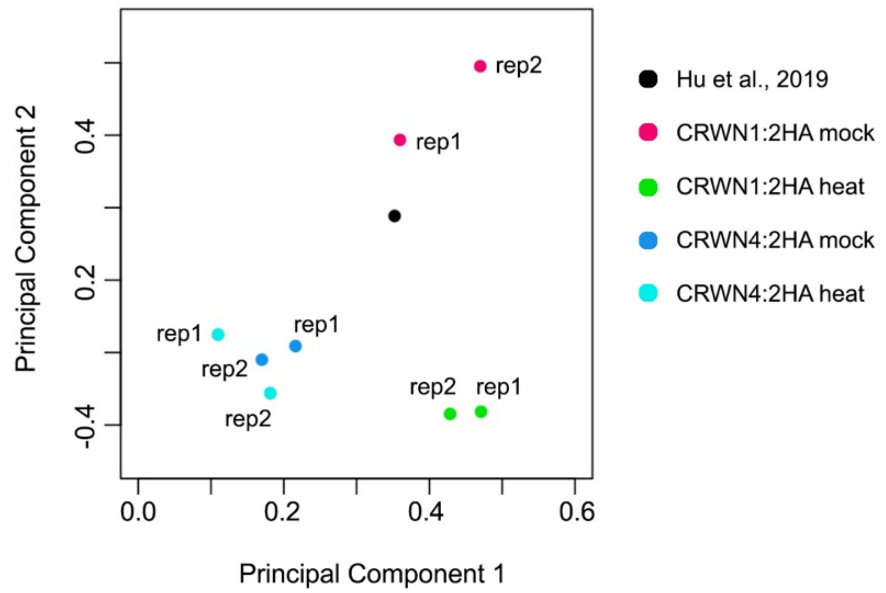


**Supplemental Figure 6. Principal component analysis of ChIP-seq samples.** Genome-wide ChIP-seq signals (normalized against input) at 20 kb window size are compared. The black dot indicates a previously published CRWN1:2HA ChIP-seq data derived from 10-day-old seedlings (Hu et al. 2019).

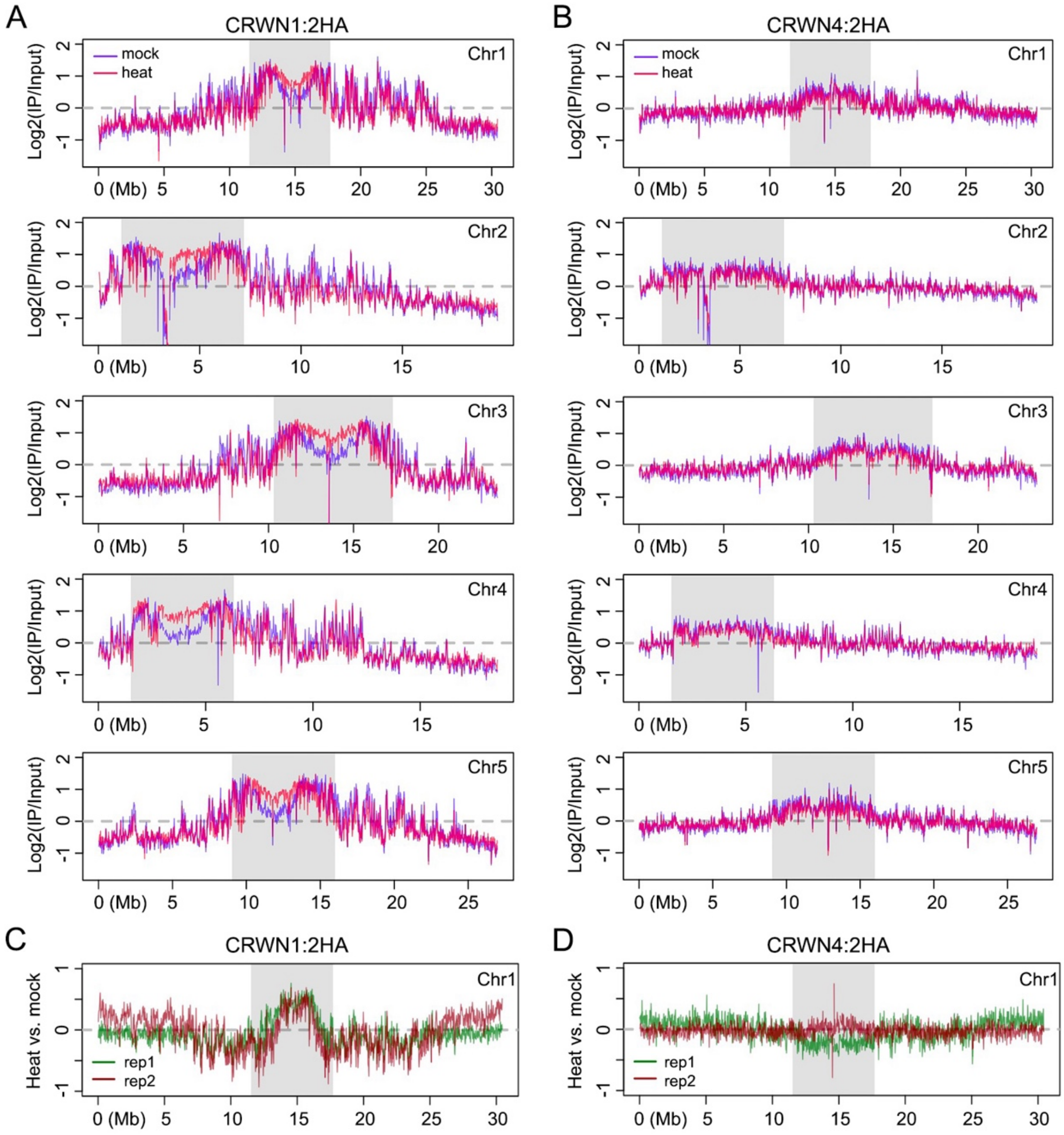


**Supplemental Figure 7. Comparison of interaction patterns between CRWNs and chromatin.**

(A, B) Genome-wide view of CRWN1:2HA (A) and CRWN4:2HA (B) ChIP-seq signals. The grey block in each plot depicts centromeric and pericentromeric regions. All the plots were generated with a 20 kb window. (C, D) Comparison of CRWN chromatin interactions with and without heat stress. The y-axis represents log2-transformed ratio of ChIP-seq signals.

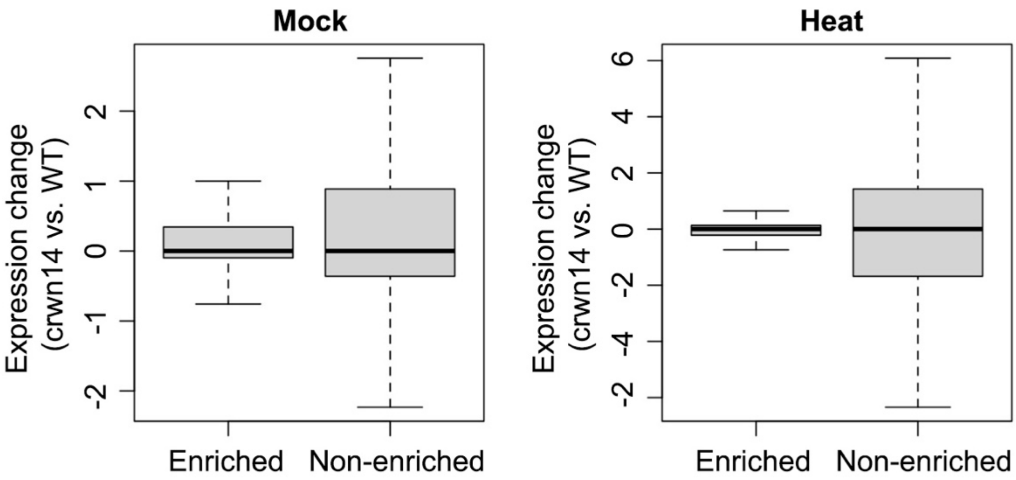


**Supplemental Figure 8. Changes in gene expression in *crwn1 crwn4* mutants.**
The boxplots show the difference of gene expression, measured in rpkm, between WT (wild-type) and crwn14 (*crwn1 crwn4*). For each growth condition, genes are separated into two groups according to CRWN1 ChIP-seq data; Mann-Whitney U test results comparing the two boxplots indicated a *p*-value larger than 0.05.

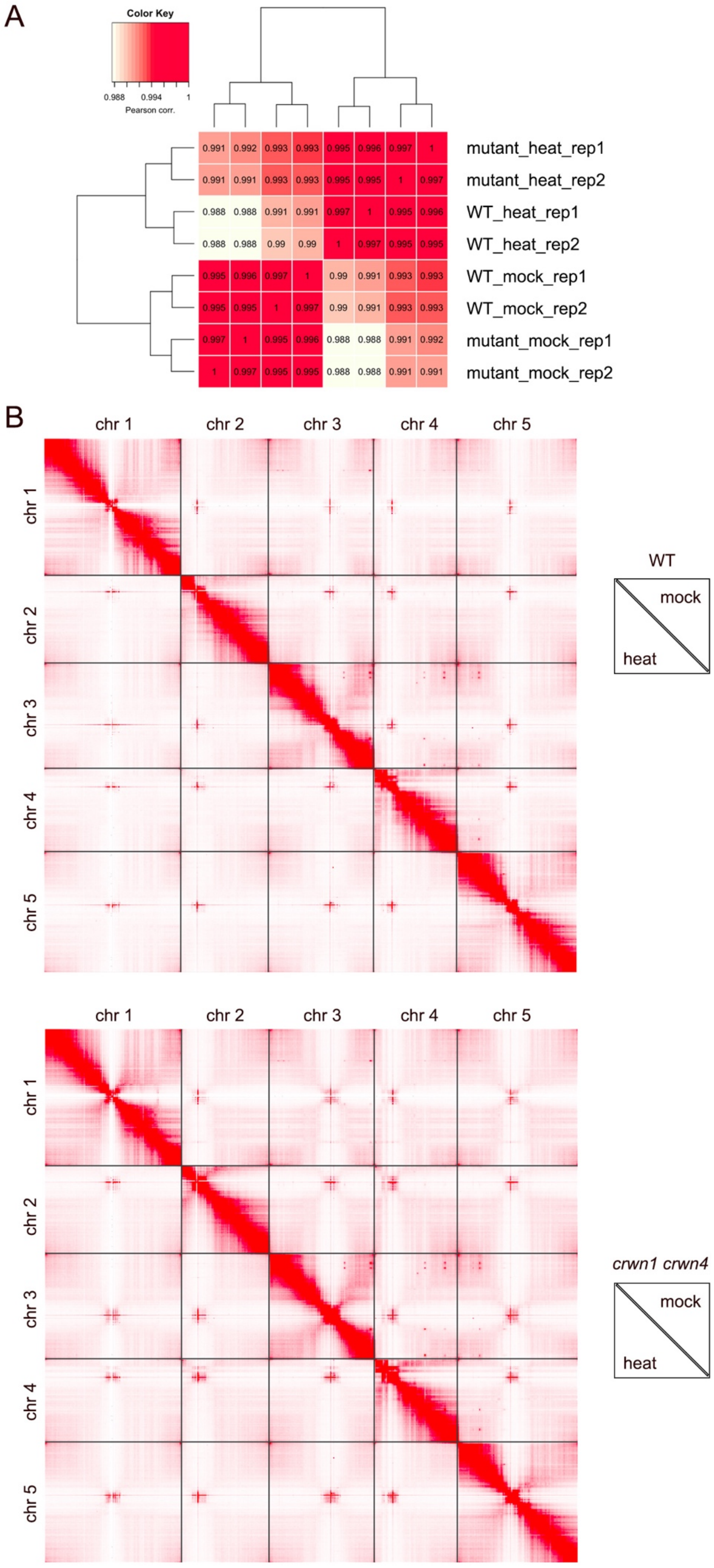
A
Color Key
0.988 0.994 1
Pearson corr.
0.991 0.992 0.993 0.993 0.995 0.996 0.997 1
0.991 0.991 0.993 0.993 0.995 0.995 1 0.997
0.988 0.988 0.991 0.991 0.997 1 0.995 0.996
0.988 0.988 0.99 0.99 1 0.997 0.995 0.995
0.995 0.996 0.997 1 0.99 0.991 0.993 0.993
0.995 0.995 1 0.997 0.99 0.991 0.993 0.993
0.997 1 0.995 0.996 0.988 0.988 0.991 0.992
1 0.997 0.995 0.995 0.988 0.988 0.991 0.991
mutant_heat_rep1
mutant_heat_rep2
WT_heat_rep1
WT_heat_rep2
WT_mock_rep1
WT_mock_rep2
mutant_mock_rep1
mutant_mock_rep2
B
chr 1
chr 2
chr 3
chr 4
chr 5
WT
mock
heat
chr 1
chr 2
chr 3
chr 4
chr 5
*crwn1 crwn4*
mock
heat

**Supplemental Figure 9. Hi-C contact maps of wild-type and *crwn1 crwn4* mutant.**
(A) Pairwise comparison of Hi-C data normalized with 20 kb bins. The dendrogram shows hierarchical clustering based on Euclidean distance. Mutant and WT stands for *crwn1 crwn4* and wild-type, respectively. Numbers in individual cells indicate Pearson correlation coefficient. (B) Genome-wide chromatin contact maps normalized with 20 kb bins.

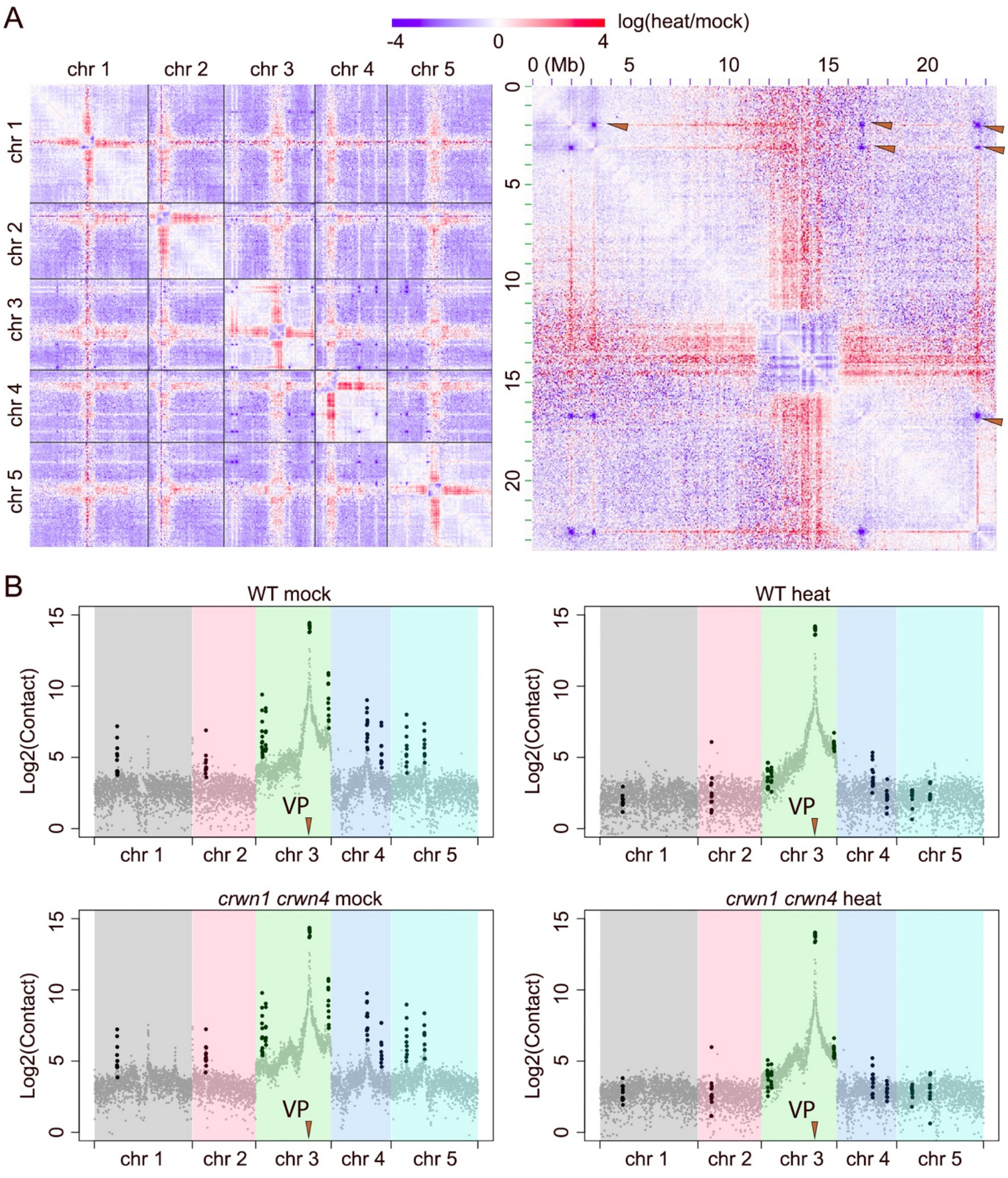


**Supplemental Figure 10. Heat stress largely weaken chromatin contact among KEE regions.**

(A) Comparison of genome-wide wild-type Hi-C contacts (left) and those at chromosome 3 (right). Colors indicate the difference of chromatin interaction strengths, expressed as the ratio of heat-stressed over mock sample. The arrowheads highlight depleted interactions between KEE regions in heat-stressed plants. (B) Contact patterns of a KEE region in chromosome 3 with the entire genome, whose location is indicated with an arrow head (VP, view point). In each panel, KEE regions (Grob et al. 2014) are highlighted with black dots.

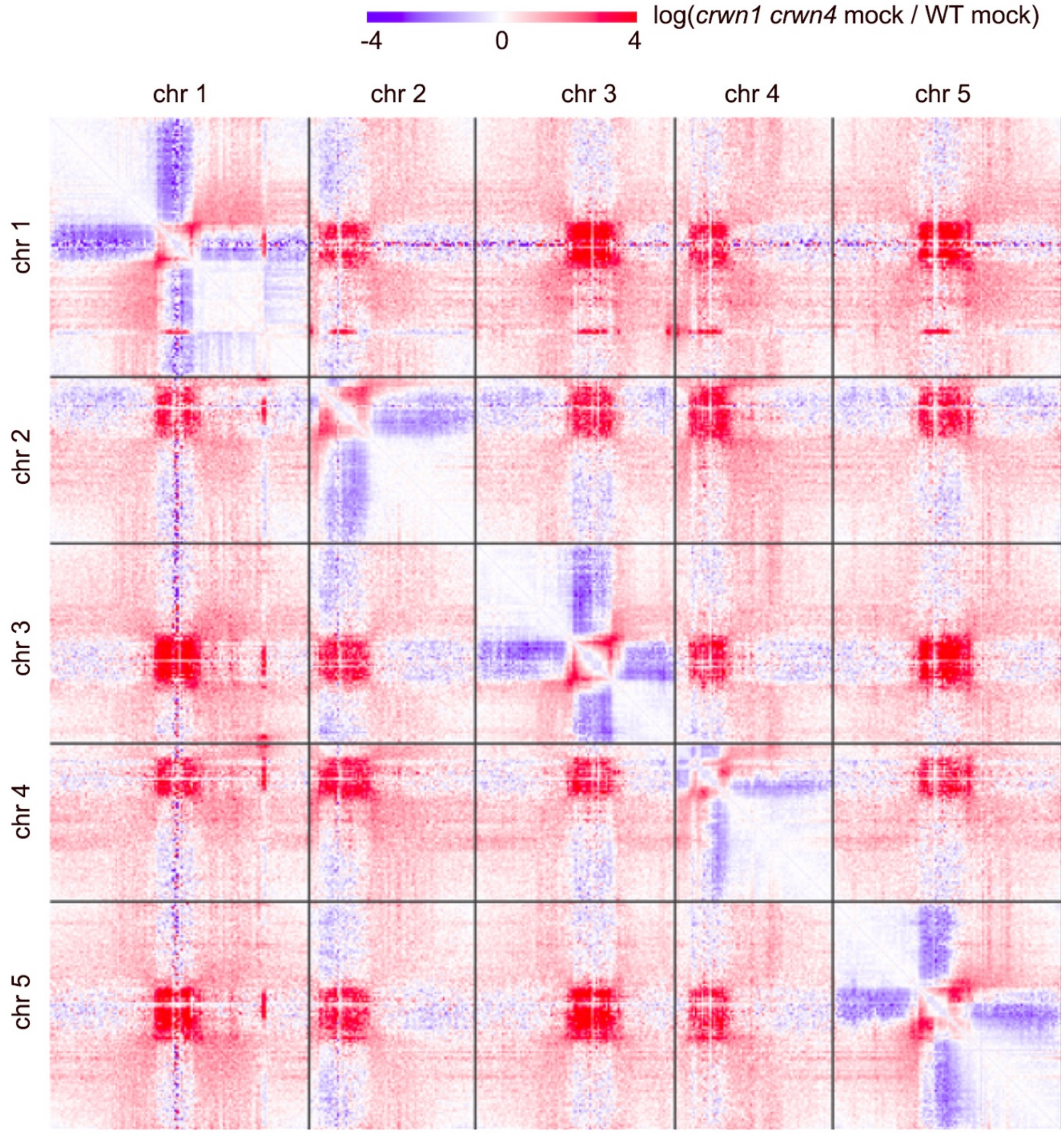


**Supplemental Figure 11. Comparison of genome-wide chromatin interaction patterns between *crwn1 crwn4* and wild-type (WT) plants.**
Colors indicate the difference of chromatin interaction strengths, expressed as the ratio of the *crwn1 crwn4* mutant over wild-type sample.

References:

Grob S, Schmid MW, Grossniklaus U. 2014. Hi-C analysis in Arabidopsis identifies the KNOT, a structure with similarities to the flamenco locus of Drosophila. *Mol Cell* **55**: 678-693.

Hu B, Wang N, Bi X, Karaaslan ES, Weber AL, Zhu W, Berendzen KW, Liu C. 2019. Plant lamin-like proteins mediate chromatin tethering at the nuclear periphery. *Genome Biol* **20**: 87.
